# Strategy Sets the Scene: Genetic architecture of linalool resistance in *Botrytis cinerea*

**DOI:** 10.64898/2026.04.05.716576

**Authors:** Melanie Madrigal, Jenna C. Moseley, Daniel J. Kliebenstein, Jordan A. Dowell

## Abstract

*Botrytis cinerea* is a necrotrophic fungal pathogen that infects thousands of plant species. During infection, these diverse plant hosts produce different specialized metabolites that can inhibit pathogen growth and shape pathogen fitness. However, the genetic architecture of pathogen resistance toward individual host defense metabolites remains poorly understood. To address this question, we exposed 83 *B. cinerea* isolates to the metabolite linalool and quantified metabolic and structural responses. Exposure revealed extensive phenotypic diversity across isolates. Population structure analysis using genome-wide SNP data showed little genetic subdivision despite diverse hosts and geographic location. Genome-wide association identified 98 genes of interest associated with membrane transport and stress response regulation. Genetic associations were stronger for morphological traits than for metabolic traits, suggesting that hyphal architecture may have a complex genetic architecture contributing to linalool resistance. Together, these results establish natural variation in linalool response and provide candidate loci for understanding how generalist pathogens respond to host-derived chemical defenses.

**Article Summary:** To understand how a generalist pathogen responds to host defenses, we asked how *Botrytis cinerea* responds to linalool, a widespread monoterpene involved in plant defense. We exposed 83 *B. cinerea* isolates to 1000 µM of linalool for 72 hours and quantified metabolic traits (growth curves and growth dynamics over time) and morphological traits (hyphal network features). Using GWA, we linked phenotypic variation to genetic variants. Results indicate substantial natural variation in linalool resistance and distinct genetic architectures across trait classes: metabolic responses are driven by a relatively small number of loci with larger effects, whereas structural/morphological responses appear more polygenic.

## Introduction

Plant-pathogen interactions are dynamic, often shaped by host and pathogen genetic factors, as well as by other complex evolutionary and ecological processes. Plants respond to pathogens through multiple defenses, including changes in metabolism, such as the induction of the jasmonic or salicylic acid pathways. These signaling pathways coordinate the production of specialized metabolites (SMs). These SMs vary in structure, abundance, and toxicity across different plant species, serving as chemical barriers that can inhibit pathogen growth^24^. Among SMs, volatile organic compounds (VOCs) can mediate information transfer between hosts and their pathogens, along with directly inhibiting pathogen growth^29^. VOCs are released in response to herbivore or pathogen attack and abiotic stressors, indicating damaged plant tissue or compromised immunity. At the same time, these VOCs can prime plants for future attacks, such that post-exposure plants mount more severe responses when they are attacked^10,15,32^. Thus, VOCs have been a major target for plant-derived pesticides due to their low molecular weight, high vapor pressure, low water solubility, and their ability to mediate resistance to multiple biological targets^59^. Due to the multifaceted effects of VOCs on pathogens and reduced environmental persistence, they are an attractive alternative to conventional synthetic pesticides^59^.

As VOC profiles vary with host identity^62^ and environment^25^ they provide rich information about host identity and condition^16^. Microorganisms may perceive these VOCs as a signal of host vulnerability. For example, the slime mold *Physarum polycephalum* exhibits chemotaxis toward/away from multiple VOCs^35^, with linalool causing *P. polycephalum* to direct its growth away from the compound. Among plant-associated habitats, such as the rhizosphere, fungi respond to plant VOCs, which can modulate fungal development and influence regulatory pathways associated with stress responses and metabolism^27^.

The generalist necrotrophic pathogen *Botrytis cinerea* provides an ideal system for investigating fungal responses to VOCs. *B. cinerea* infects thousands of hosts with few genetic indicators of host specificity^6,13^ despite evidence that gene expression responses can be host specific^45^. As such, virulence in *B. cinerea* is polygenic, with small-effect loci contributing to infection success rather than a few major-effect loci^6,13,54^. Moreover, it relies on a multitude of broad-spectrum virulence strategies, with genome-wide association studies demonstrating that variation in virulence genes is distributed across the genome and shows little enrichment for specific pathways or compartments^5,6,13^. As a necrotrophic pathogen, *B. cinerea* relies not only on host tissue degradation but also on its capacity to overcome host defenses through a variety of mechanisms, including enzymatic detoxification and transporter-mediated efflux^63,66^. Further, the sheer diversity of hosts suggests that *B. cinerea* individuals may be exposed to a wide range of lineage-specific and general VOCs, necessitating strategies to deal with an array of complex SMs and VOCs alike.

Among plant VOCs, linalool is found in over 200 monocotyledonous and dicotyledonous plant species, belonging to over 50% of plant families^4^. Linalool is an acyclic monoterpene alcohol that is produced through the methylerythritol phosphate (MEP) pathway, where geranyl diphosphate (GPP) serves as its immediate precursor and is catalyzed by linalool synthases, members of the large terpene synthase gene family, and is responsible for the production of other monoterpenes. Reported concentrations of linalool range from 0.01 ppm to over 0.6 ppm in fruit tissue and from 1.6 ppm up to 20000 ppm in leaves.

As a VOC, linalool plays several roles, from attraction to defense. Linalool is a major constituent of the floral scent of many species and is widely used to attract pollinators, which rely on sensory cues to find suitable hosts^31^, and as a cue for nearby plants, informing them of environmental conditions^27^. Linalool also acts as a direct defense mechanism against pathogens. For example, in *Pseudomonas fluorescens,* linalool reduces membrane potential, increases cellular leakage, and decreases protein content^28^. Similar modes of action have been observed in fungal pathogens, including *Candida*^33^, and *B. cinerea*. Specifically, Xu et al. reported that when exposed to linalool, a *B. cinerea* strain downregulates ergosterol biosynthesis genes and upregulates genes encoding CYP450 enzymes, a large and diverse family of enzymes involved in xenobiotic detoxification, specialized metabolism, and lipid modification ^65^. Reduced ergosterol biosynthesis compromises plasma membrane integrity, disrupting membrane-associated signaling processes and membrane structure, as ergosterol is a vital component of the fungal plasma membrane. In addition to enzymatic detoxification, other studies have identified major facilitator superfamily transporters (MFS) as key components of resistance and virulence in *B. cinerea.* These transporters mediate the efflux of toxic compounds from the cell, reducing the accumulation of plant defensive SMs or other pesticides. Beyond its direct effects on pathogens, linalool also acts indirectly by stimulating plant-host hormonal pathways. When *Orzo sativa* was exposed to linalool, it upregulated defense genes and demonstrated enhanced resistance to the bacterial pathogen *Xanthomonas oryzae*^58^.

While there is support for the direct and indirect effects of linalool on plant pathogen interactions, most studies focus on a single or a few isolates^53,65^. Although these approaches provide valuable mechanistic insight, they capture only an individual genotype’s response and do not necessarily reflect how individuals across a genetically diverse population respond or the strategies they employ. Given the polygenic architecture of virulence in *B. cinerea* and high effective population size^13^, this suggests that candidate genes for linalool resistance validated in the model isolate of *B. cinerea* (BO5.10)^65^ may not capture the full diversity of resistance mechanisms employed across the species. While genome-wide association studies (GWAS) have been successfully applied to virulence traits, few studies have examined the genetic architecture of *B. cinerea’s* responses to common specialized metabolites at the population level^18^. To address this gap, we exposed a population *of B. cinerea* isolates to linalool to assess population-level variation in resistance and to investigate underlying trait genetic architecture.

## MATERIALS AND METHODS

### Botrytis Collection

We leveraged 83 isolates of the haploid *B. cinerea* association mapping population previously described^5,7,13^.Isolates were obtained from more than 14 different plant species, of which 90% were isolated in California, and 10% were worldwide strains regularly used for functional genomic studies^3,14,45^. Within this population, there is little evidence of host specialization or local population structure indicating high global gene flow^19,39,51,55^, as Casey et al. found Fst values of 0.03 between strains isolated in Europe and California, and 0.02 among isolates from different plant host species^13^. All strains used in this study have been previously whole-genome sequenced to an average of 164-fold coverage^5,7^ and aligned to the genome assembly of B05.10 ASM83294v1^60^, resulting in 217,749 SNPs across the approximately 50-Mb genome, demonstrating substantial standing genetic variation within the population and providing dense genome-wide marker coverage for association analyses^5,7^.

### Linalool Exposure

The isolates used in this study were grown for 2 weeks on oatmeal agar plates. Conidia were collected from full grown plates using sterile water, counted using an automated spore counter pipeline(https://github.com/melaniemadrigal07/Post.ImageProcessing) and diluted to a final concentration of 10 spores/µL in a gamborg B5 (Sigma Aldrich, St. Louis MO) supplemented with 41 g/L fructose (Thermo Fisher Scientific, Waltham, Massachusetts) 26.65 g/L glucose (Sigma Aldrich, St. Louis MO), 1.03 g/L sucrose (Fisher Scientific, Pittsburgh, PA), with pH adjusted to 6.5 using 0.01 KOH. To prevent spore germination, spores were kept on ice from collection to inoculation. 10 µL of the spore suspension (10 spores/µL) were inoculated into a 96-well plate with 70 µL of liquid Gamborg B5, 10 µL of 1000 µM Linalool (0.045% DMSO (Sigma Aldrich, St. Louis, MO), and 10 µL of OZ blue cell viability (OZBiosciences, San Diego, CA), in an 8-fold replication. The 1000 µM linalool concentration was selected based on its biological relevance, as reported concentrations range from 0.06 µM to 130,000 µM in fruit tissues^22^. In line with this range, Xu et al. demonstrated minimum inhibitory concentration (MIC; the lowest dose that inhibited spore germination) and fungicidal concentrations (MFC; the lowest dose that inhibited mycelial growth) for *B. cinerea* (MIC=20.95 µM; MFC=2,095 µM)^65^, placing 1000 µM within a physiologically meaningful window that is consistent with host-derived exposure and below fungicidal thresholds.

The OZ blue cell viability assay relies on mitochondrial metabolic activity in viable cells: the blue-redox dye is taken up and reduced in the mitochondria, resulting in a detectable color change. Plates were imaged every 4 hours for 72 hours for a total of 20 time points using a Cytation Gen5 Plate (Agilent, Santa Clara, CA) coupled to a Biospa (Agilent, Santa Clara, CA). Quantitative colorimetric data were extracted from the time-lapse images for subsequent analysis of respiratory rates.

### Image Analysis

The Cytation Gen5 generated approximately 336,000 images, each exported as a separate red, green, and blue (RGB) channel. To organize data for analysis, an ImageJ macro was used to group and compile images by time point for each plate, for downstream analysis. Using these images, the RGB channels were converted to the L*a*b* color space, and values were extracted using a custom Python script https://github.com/melaniemadrigal07/GWASLIN). With the L* value measuring the shift from light to dark, when L = 0, it indicates a dark appearance, and L = 100 indicates a bright appearance. The a* value quantifies the shift from green to red, with negative a* indicating green and positive a* indicating a strong red color. The b* value is a proxy for the shift from blue to yellow, with a negative b* indicating a strong blue component and a positive b* indicating a strong yellow component.

To accurately extract values from the images, a 10% L-value threshold was applied to isolate hyphal growth in each well. We then measured values external to hyphae, thereby enabling quantification of color changes in the media associated with fungal metabolic activity.

After extracting the color values, background correction was performed by subtracting the average L*, a*, and b* values from blank wells (media + dye without spores) from those of experimental wells at each time point.

### Metabolic Activity Quantification

Generated L*, a*, b* values were combined using the following formula to obtain an ΔE Value: 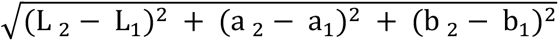, ΔE serves as a proxy for color change relative to the first time point. This metric was used to generate a curve representing color change over time. To capture early metabolic activity, we focused on the early germination phase by using timepoints one to five (up to 20 hours post-inoculation) for downstream analyses. During this 20 hpi window, spores initiate germination and begin forming penetrative structures^13,17^ (e.g., appressoria, infection cushions). In addition to ΔE, b* values were analyzed independently, as they reflect the reduction of the resazurin dye and provide a more specific measure of redox-driven metabolic activity.

Several phenotypes were extracted from the ΔE curve (Supplementary Table 1), area under the curve, and maximum ΔE. The area under the curve represents the magnitude of the color change at the early time points. Max ΔE is the value at which isolates reached their peak color change. Similarly to the phenotypes extracted from ΔE, the same was done for the b* value, which represents the shift along the blue-yellow axis, with positive values indicating yellow and negative values indicating blue. The color change range was incorporated as a phenotype to represent metabolic variation in response to linalool.

To examine the foraging strategies of the strains, hyphal network images were skeletonized using SkelPy and subsequently processed with SkelpyR to remove skeletonization artifacts^41^.Network-level traits were extracted, including edge lengths, total hyphal length, number of tips, number of branches, tip fraction, and branch fraction. Total hyphal length and tip number provide measures of overall growth and active extension, while tip and branch fraction capture how growth is allocated between exploratory hyphal tips and network consolidation, and edge length provides information on the spatial scale of individual hyphal segments.

Prior to multivariate analyses, trait distributions and pairwise relationships were evaluated using frequency histograms, scatterplots, and Pearson correlation analyses. Scatterplot matrices were used to visualize distribution of each trait and pairwise correlations among traits (Supplementary Figure S1). A principal component analysis (PCA) was used to summarize multivariate patterns of phenotypic variation among *B. cinerea* isolates. PCA was conducted separately for structural hyphal network traits and metabolic traits, using the same framework for both datasets. PCAs were performed on centered and scaled trait matrices using singular value decomposition, excluding traits with zero variance. K-means clustering was subsequently applied to isolate eigenvectors corresponding to the first two principal components. The number of clusters was set to k=2 based on inspection of the within-cluster sum of squares and the elbow method. Clustering was used to summarize the major axes of phenotypic variation rather than to define discrete phenotypic classes.

### Genome-wide Association Study

To estimate the genetic architecture of resistance to linalool in *B. cinerea,* we performed genome-wide association using GEMMA’s univariate linear mixed model^69^. Genotype data of our population was derived from the SNP dataset generated by Atwell^7^ and was converted to PLINK binary format for downstream analyses. Kinship relatedness among isolates were accounted for using a centered genotype matrix in GEMMA. This approach was based on the model of Zhang et al.^67^and further adapted by Dowell et al.^21^ In this framework, Single Nucleotide Polymorphisms (SNPs) with a minor allele frequency (MAF) of > 0.20 and < 10% missing data were retained, yielding 271,495 SNPs for analyses. Population structure was estimated independently using a principal component analysis (PCA) implemented in PLINKv1.90. Prior to running a PCA, linkage disequilibrium (LD) pruning was calculated by looking at a window size of 50 kb, 10 steps and a r^2^ threshold of 0.1 to reduce influences of highly correlated SNPs. Previous studies have identified that linkage disequilibrium decay occurs in small regions (<1,000).^8^ Following the PCA, the elbow method was used to determine the optimal number of components to retain for downstream model analysis. Two different association models were run per trait. A kinship (K) model and a kinship plus population structure in which the first three principal components were included as co-variates in the (P+K) model to capture population structure. To assess mixed linear model performance, quantile-quantile (q-q plots) plots were used. In q-q plots, the observed p-value is plotted against expected probability distribution to identify where the model may have produced higher number of significant results due to chance (Supplementary Figure S2). Due to low linkage disequilibrium in *B. cinerea*^5^, association significance was assessed using a Bonferroni correction with the family-wise error rate set at 𝛼 = 0.05 divided by the number of SNPs tested.

Following GWA, we assessed linkage disequilibrium (LD) among significantly associated SNPs using PLINK^47^. Pairwise LD (R^2^) was calculated for SNPs within 10 kb windows centered on significant SNPs to assess regional associations. SNP effect sizes and standard error were calculated as the beta divided by the trait range, as *B. cinerea* is haploid^43^. Relative effect sizes were summed across all associated regions for each trait to estimate the relative effect size of the associated genomic regions. Differences in the distribution of effect sizes between metabolic and structural traits were assessed using a Wilcoxon rank-sum test in R. Regions associated with multiple traits were considered colocalized. Gene functional annotations, when available, were extracted from the fungal genomic resources^9^. SNPs were further classified by genomic location (coding or non-coding) and mutation type (e.g., synonymous, missense, or nonsense).

To prioritize associated loci for downstream interpretation, we identified lead SNPs exceeding the genome-wide significance thresholds (P < 0.05/m, where m is the number of SNPs tested) and defined associated regions using local linkage disequilibrium patterns around each lead SNP (pairwise R^2^ within 10 kb). Among significant regions, loci were ranked by lead SNP P-value, standardized effect magnitude, and evidence of colocalization across traits. Candidate genes were defined as those overlapping each region or as the nearest to the lead SNP. For candidate SNPs in protein-coding genes, we assessed codon bias among synonymous SNPs by reconstructing reference and alternate codons for each SNP (using SNP context, CDS position, and gene strand) using a custom R script (https://github.com/melaniemadrigal07/GWASLIN). Usage was then compared with the genome-wide codon usage frequencies of B05.10’s reference genome^57^. All analyses were conducted within the R environment^48^ (R Core Team 2025, version 4.5.3).

### Molecular Docking

To explore the potential role of identified variants that directly interact with linalool, rather than via indirect effects on gene expression or the overall cellular environment, we used an in-silico protein-ligand binding approach. For each protein of interest, we used D-I-TASSER^50,68^ with default settings to predict its structure. D-I-TASSER was selected due to its demonstrated performance in CASP benchmarking. I-T Model confidence was assessed using the reported Cscore (a confidence score that estimates the quality of predicted models) and the estimated TMscore (a metric for assessing the topological similarity of protein structures), and the top-ranked model was used for downstream analyses. To assess the interaction of linalool with predicted protein transporters, we used CaverDock^42^ and AutoDock Vina to estimate the binding channel for simulation, employing a blind docking approach when proteins lack previously identified and validated active sites. The protein Bcin07g05680 was selected among genes with significantly associated SNPs as a candidate for docking analysis because it encodes an MFS transporter and lacks an identified active site. Given its association with metabolic traits, we used a blind docking approach to identify potential binding sites for linalool.

## RESULTS

### Metabolic activity

Across *B. cinerea* isolates, the Area Under the Curve Δ𝐸 varied from 3.57 to 109 among samples, with an average of 17.99 (±2.2 𝑆𝐸, supplementary Table 2). The slope from timepoint 5 (20 hours post inoculation) varied from -0.01 to 2.37, with an average of 0.08 (±0.01 𝑆𝐸, Supplementary Table 2). For traits related to the shift along the b* axis, the maximum blue varied from -1.15 to 24.69, with an average of 4.66 (±0.65𝑆𝐸, Supplementary Table 2). The slope of b* ranged from -0.86 to 6.40 among samples, with an average of 0.92 (±0.17 𝑆𝐸, Supplementary Table 2). Area under the curve along the b axis varied from -6.47 to 79.89 among samples, with an average of 11 (±1.61𝑆𝐸, Supplementary Table 2).

Principal component analysis (PCA) of the ΔE trajectories and blue-axis traits separated isolates along PC1 and PC2. The PC1 axis accounted for 76.3% of the total variance and captured the magnitude of the metabolic response, whereas PC2 accounted for 12.6% of the variance and described differences in response timing and the shape of the metabolic activity trajectory (Figure 1B).

**Figure 1.**
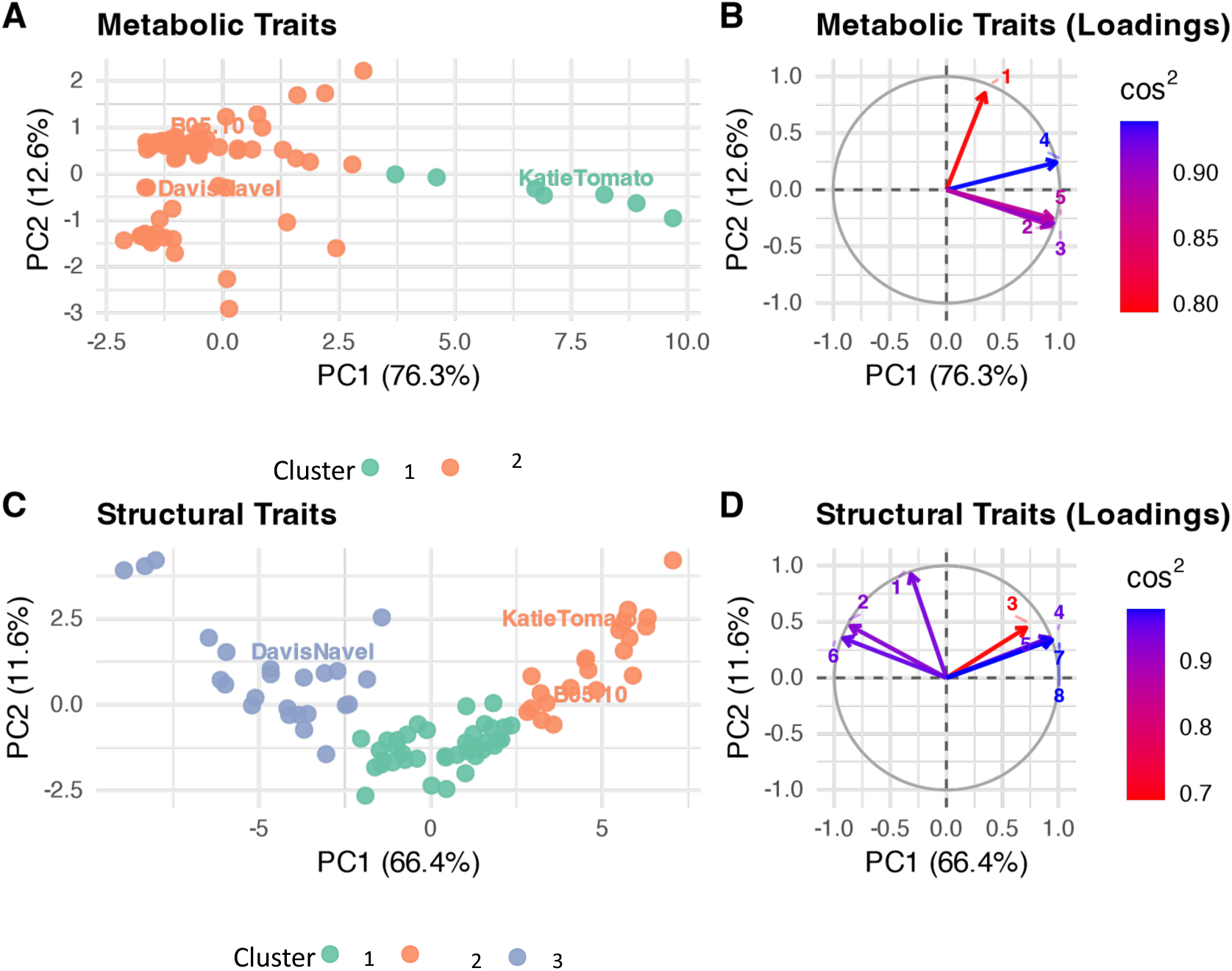
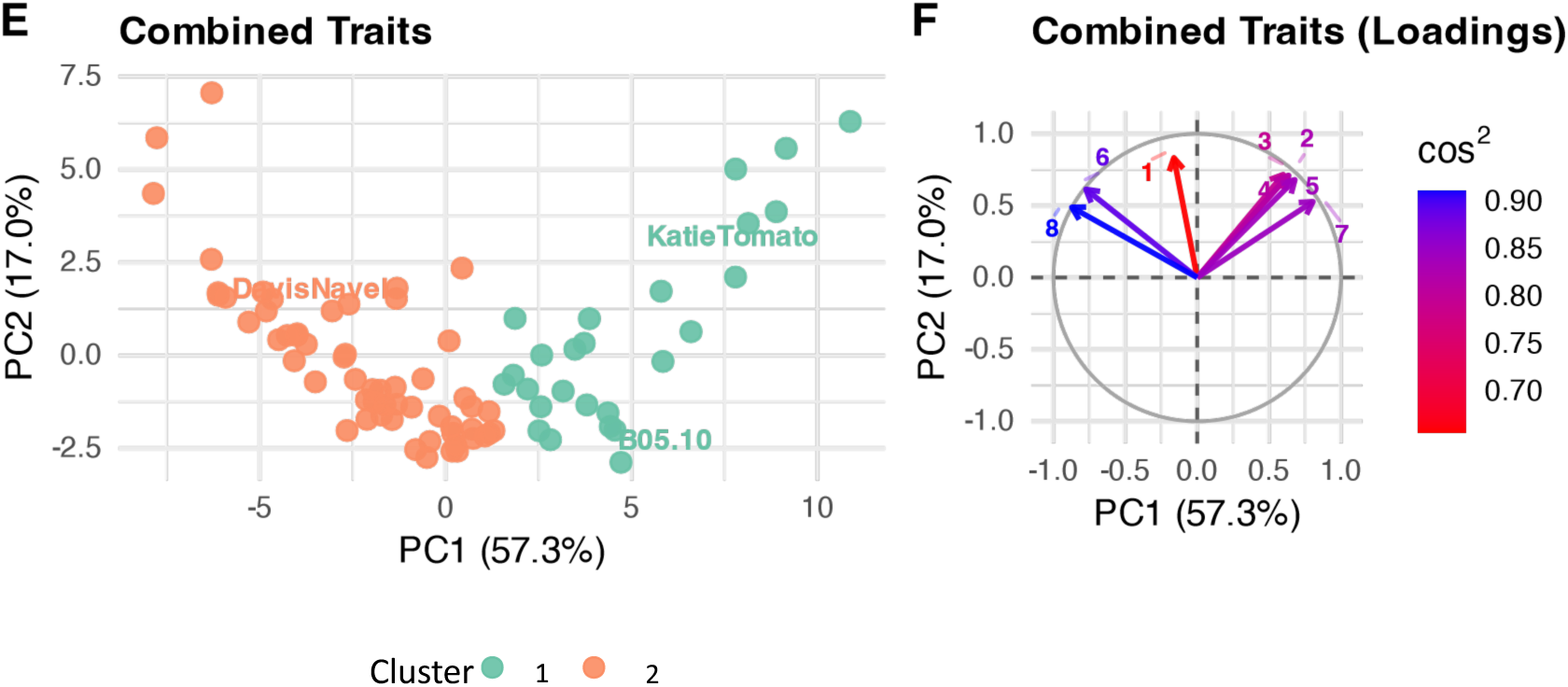
Metabolic and structural trait variation across isolates. (A, C, E): PCA score plots of *B. cinerea* isolates projected onto PC1 and PC2 for metabolic (A), structural (C), and combined (E) trait datasets. Points are colored by k-means cluster assignment (k=3 for structural, 2 for metabolic, and combined), selected isolates are labeled for reference, including B05.10 (reference and generalist isolate), KatieTomato (highly virulent isolate), DavisNavel (low virulent isolate). (B, D, F) Corresponding loading plots displaying the top contributing traits for 51each dataset. Arrows represent variable loadings on PC1 and PC2. In (B), Traits in numerical order: 1, time point of maximum blue axis shift; 2, maximum at timepoint 5 (20 hpi); 3, area under the curve at timepoint 5 (20 hpi); 4 slope of blue axis shift at timepoint 5 (20 hpi); 5 slope of ΔE at timepoint 5. (D) Traits in numerical order: 1, tip fraction residuals; 2, weighted tip fractions; 3, area under the curve of fractal dimension across 72 hours; 4, total hyphal length; 5, mean total hyphal length; 6, median edge length; 7, number of hyphal tips; 8, number of hyphal nodes. (F) 1, tip fraction residuals; 2, maximum blue axis shift; 3, range of blue axis; 4 slope of blue axis shift at timepoint 5 (20 hpi); 5, area under the curve of blue axis at timepoint 5 (20 hpi); 6, tip fraction weighted; 7, area under the curve of ΔEat timepoint 5 (20 hpi) 8, median edge length.

### Hyphal network traits

Structural traits of *B. cinerea* isolates, such as fractal dimension, varied from 0.19 to 1.44, with an average of 0.72 (±0.02 𝑆𝐸,Supplementary Table 2). The slope of the fractal dimension varied from -0.11 to 0.19, with an average of 0.05 (±0.006 𝑆𝐸, Supplementary Table S1). Area under the curve of fractal dimension ranged from 1.01 to 8.73 (±0.18 𝑆𝐸,Supplementary Table 2). Cumulative total hyphal length across the time course ranged from 778 to 144,171 µm with an average of 37,519 µm (±0.02𝑆𝐸, Supplementary Table 2). Architectural ratios such as tip fraction ranged from 0.702 to 0.937, with an average of 0.78 (±0.02 𝑆𝐸, Supplementary Table 2). The number of branch points (locations where hyphae bifurcate) ranged from 0.37 to 403, with an average of 122 (±10.7 𝑆𝐸, Supplementary Table 2). The number of tips (proxy for active regions of the hyphal region) ranged from 4.6 to 863, with an average of 294 (±20.3 𝑆𝐸, Supplementary Table 2). Edges representing the connectivity of the hyphal network ranged from 2.99 to 1279, with an average of 404 (± 67.𝑆𝐸, Supplementary Table 2).

A PCA of structural traits revealed three distinct clusters among *B. cinerea* (Figure 1C). PC1 accounts for 69.7% of the variance and is largely driven by connectivity-related traits (high AUC, fractal dimension, tips, and edges). PC2 accounts for 12.1% of the variance and is driven by growth metrics (fractal dimension, slope, and fragmentation index). A PCA of both structural and metabolic traits revealed a similar pattern; PC1 explained 57.3% of the total variance, and PC2 explained 17.0% (Figure 1C). PC1 was primarily associated with structural traits such as tips and edges. PC2 captured secondary variation, dominated by metabolic color-shift dynamics along the b* axis (Figure 1D).

### Population Structure

The principal component analysis ran on linkage disequilibrium genome wide pruned SNP data. The first three principal components were evaluated to assess population structure within our collection (Supplemental Figure 2). The first three principal components capture 30% of the variation Points were colored by host association and shaped according to geographic origin. Across our population, no clear population stratification by host or geographic location was found (Supplemental Figure 2A-2C) which coincides with previous studies^13^. If strong population stratification was present, we would expect to see isolates collected from same hosts or geographic origin cluster together within the PCA space (Supplemental Figure 2).

**Figure 2.**
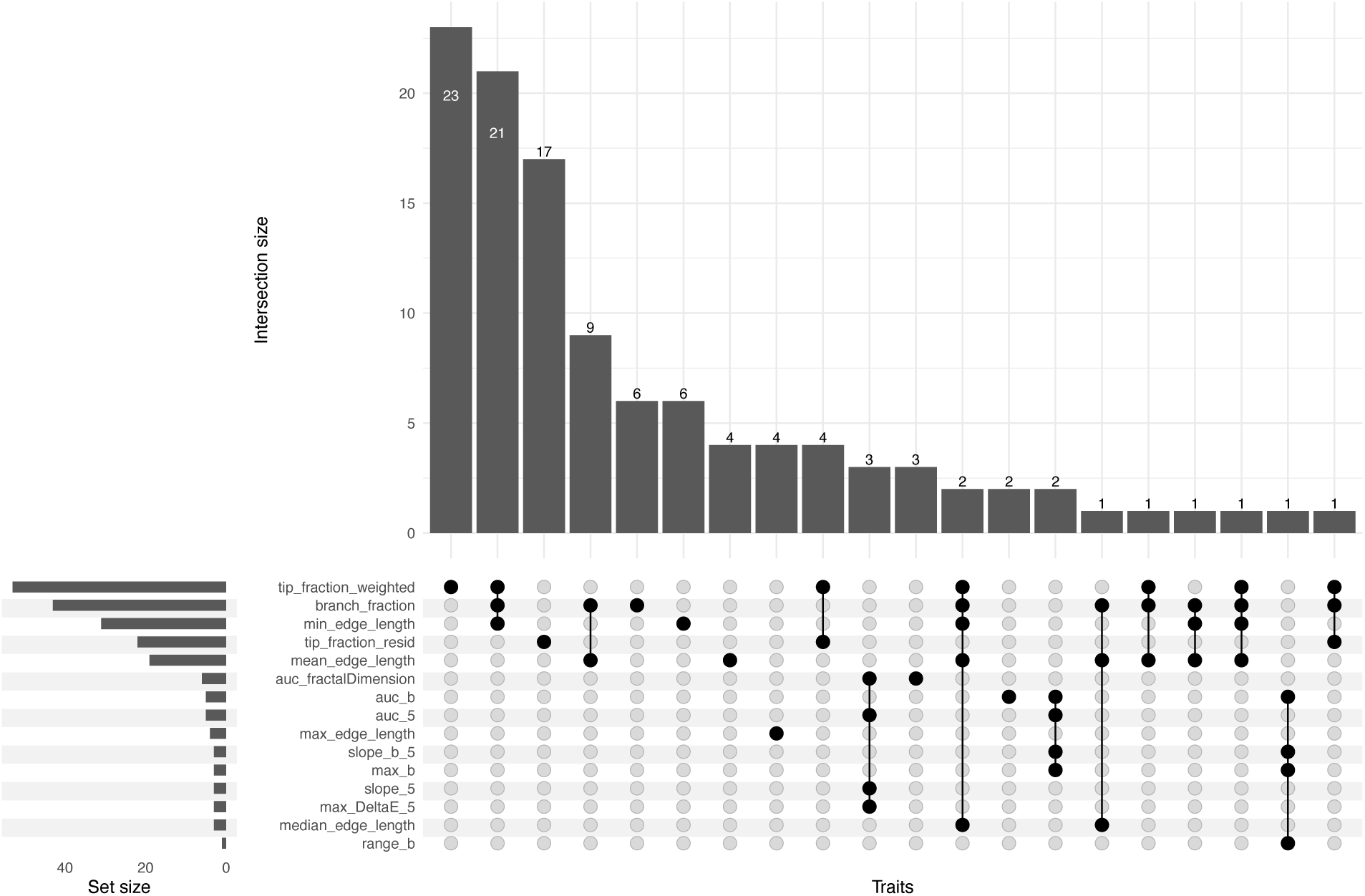
Overlap of significant SNPs and trait associations across both the kinship (K) and the population structure and kinship model (P+K). Upset plot showing the intersection. of significant SNPs across measured traits. Vertical bars represent the number of SNPs shared among specific combination of traits (intersection size). Horizontal bars indicate the total number of SNPs associated with each individual trait (set size). Connected black dots denote traits included in each intersection.

### Comparison of Mixed Models

For all traits, 2 mixed models were considered: kinship (K) and population structure, as measured by principal component analysis (P+K). Quantile-Quantile (q-q) plots of observed P values versus expected P values for each model were used to assess model accuracy (Supplementary Figure 3). The distributions of observed P values in both the kinship (K) and principal component analysis (P+K) models were similar (Supplementary Figure S3-5)., across the b* trait range (represents the variation of blue axis in the LAB axis) models exhibited systematic under inflation (Supplementary Figure 3f). While the slope of fractal dimension had high systemic over inflation. Across all traits assessed both models shared 74 SNPs, with 24 unique to the K model and 14 unique to the P+K model (Supplementary Figure S3-4). Given the limited population structure (Supplementary Figure S2) detected within the panel was minimal, the P+K model appeared more overly conservative reducing the power to detect associations. Therefore, the K model K model was used as the primary model considered for marker examination.

### Genetic architecture of resistance

Genome-wide association analyses using the traits as phenotypes identified 172 significant SNP-trait associations and 98 unique SNPs in the K model (Figure 2, Supplementary Table 3). Trait associations were most frequent on chromosomes 4 (n=32),2 (n=27), 7 (n=24), and 8 (n=23). Significant SNPs were distributed across the genome, with the greatest number on chromosome 2 (n=14; 10 non-coding,4 coding), chromosome 8 (n=13; 12 noncoding,1 coding), chromosomes 4 (n=11; 3 non-coding, 8 coding) and 13 (n=11 noncoding), all remaining chromosomes contained 8 or fewer significant SNPs (Figure 2, Supplementary Table 3). Of the 98 unique SNPs, 75 were noncoding, 16 were coding-synonymous, and 7 were coding-missense variants. Several significant SNPs were mapped within or near four genes on chromosome 7 (*Bcin07g05680*, *Bcin07g05230, Bcin07g05240, and Bcin07g05210*), two distributed across chromosomes 2 (*Bcin02g09200 and Bcin02g05460*) and 15 (Bcin15g00890), and 3 (*Bcin03g06600*). Although multiple SNPs supported individual loci, overlap among traits was limited (Figure 2). When present, trait overlap suggests that a shared locus may contribute to multiple aspects of the linalool response. In the P+K model, 14 SNPs associated with structural traits were not detected in the kinship model (Supplementary Figure 4). However, both models shared 72 significant SNPs, suggesting overall agreement in the major association signals (Supplementary Figure 4). Additional SNPs identified in the P+K model are associated with structural traits that were not detected in the kinship model. These additional associations may reflect loci influenced by subtle population structure, suggesting that some components of the response to linalool vary across subpopulations within the *B. cinerea* panel. Our *B. cinerea* panel consists of individuals collected from different hosts. Variation within host’s chemical diversity and production of linalool may contribute to these differences in linalool-resistance associated alleles and response strategies. Similar patterns have been explored in a fungicide resistance population, in which strong genetic differentiation among resistance isolates was present despite little population structure across hosts, growing cycle, or mating type.^38^

Multi-trait associations were rare, with most SNPs associated with a single phenotype, suggesting that most SNPs contribute to specific components of the linalool response rather than broadly affecting multiple traits. Here, a pleiotropy is defined as a SNP associated with multiple phenotypes. Across metabolic-activity traits (Δ𝐸 and b features), several traits shared associations within*/*near *Bcin07g05680 (*MAF=0.212), which encodes a major facilitator superfamily transporter. Notably, most SNPs within this locus were synonymous mutations. Metabolic traits exhibited slightly larger standardized absolute SNP effect magnitudes than structural traits (mean = 0.14 vs 0.13; Figure 3A). However, structural traits included substantially more unique SNPs (94 vs 7) and significant SNP-trait associations (15 vs 21) (Figure 3B).

**Figure 3.**
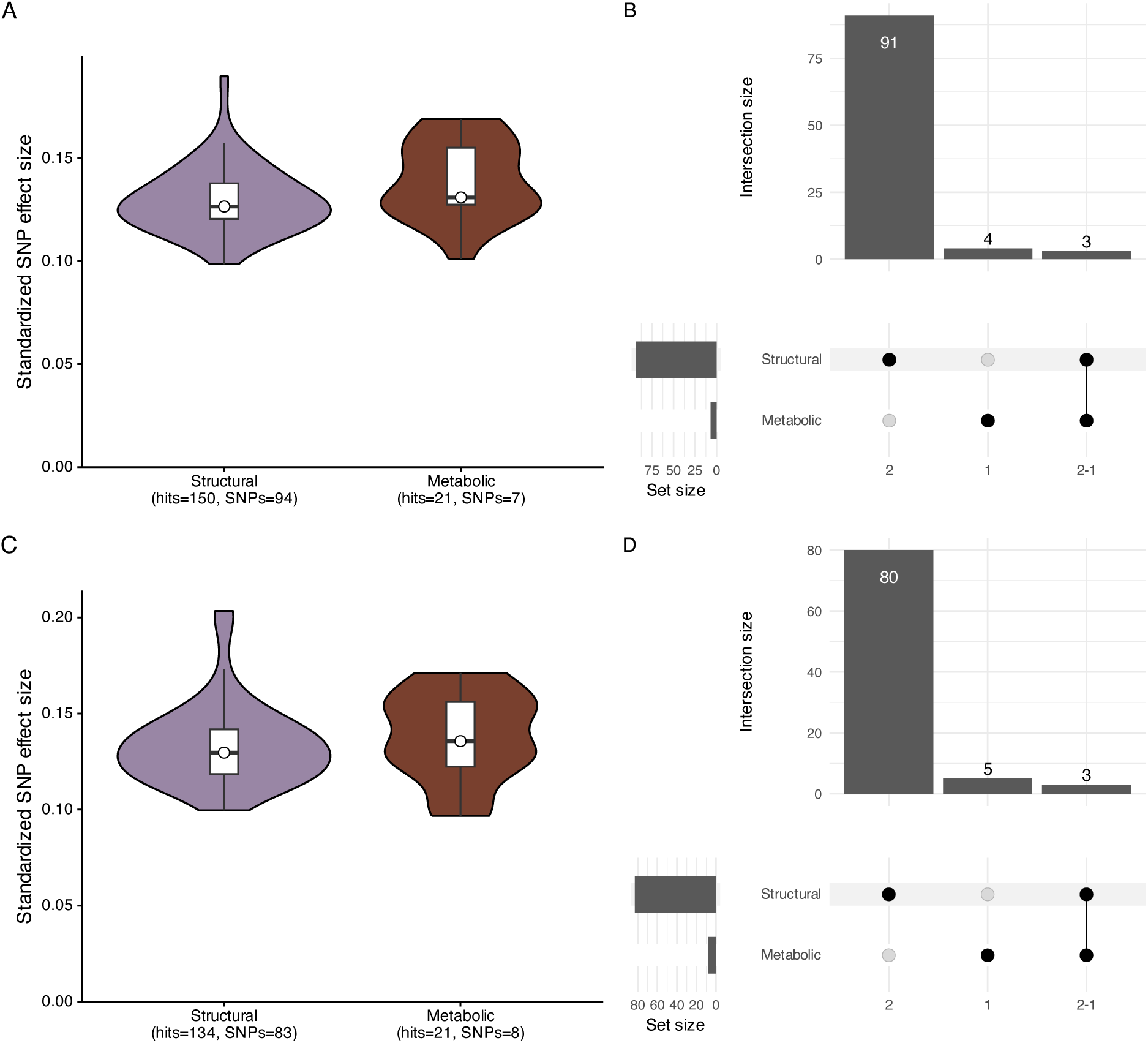
Distribution and overlap of significant SNP associations across structural and metabolic trait classes. (A,C) Distribution of standardized SNP effect sizes for structural and metabolic traits in the Kinship (K) model (A) and the population structure plus kinship (P+K) model (C). Violins represent the full distribution of absolute effect sizes across significant SNP-trait associations, and boxplots indicate the interquartile range and median. Sample sizes (total significant associations and unique SNPs are shown below each category. Hits refer to significant SNP-trait associations, while SNP counts represent unique variants after collapsing repeated associations across traits. (Wilcox rank sum test=968, P=0.0043 in the Kinship model). (B,D) Upset plot of the Kinship (K) model and the population structure plus kinship (P+K) model showing the number of unique SNPs associated with structural traits, metabolic traits, or both. Vertical bars represent interaction sizes, and the matrix below indicates the trait class for each intersections. Structural SNPs account for the most significant associations, with limited overlap between structural and metabolic traits.

Of the SNPs identified, 78 mapped to non-coding regions and 23 to coding regions. Within coding regions, 16 were synonymous, and 7 were missense variants (Figure 4A). 6 of the missense variants mapped to Bcin04g02170, a protein kinase domain-containing protein, while one variant mapped to the transcription factor Bcifm1. Within Bcin04g02170, three substitutions were altered amino acid charge, and two altered amino acid size class (Supplementary Table 3). Notably, one substitution changed arginine to glycine, thereby shifting from a positively charged residue to a nonpolar residue. Arginine is larger and can maintain its charge, allowing stronger hydrogen and electrostatic bonding, while glycine is smaller, uncharged, and has a minimal hydrogen side chain, which could increase flexibility and alter glycine’s substrate specificity. Previous work on protein kinases has shown that hydrostatic and electrostatic balance is a major determinant of kinase-substrate specificity, and that residues outside the active site can influence substrate specificity through effects on protein structure and electrostatic environments^36^. The final missense variant mapped to the translation initiation factor Bcifm1, which changed alanine to valine. Although both amino acids are nonpolar, valine is bulkier due to its branched side chain. This increase in steric volume could subtly alter local packing, protein stability, or interaction surfaces^26^.

**Figure 4.**
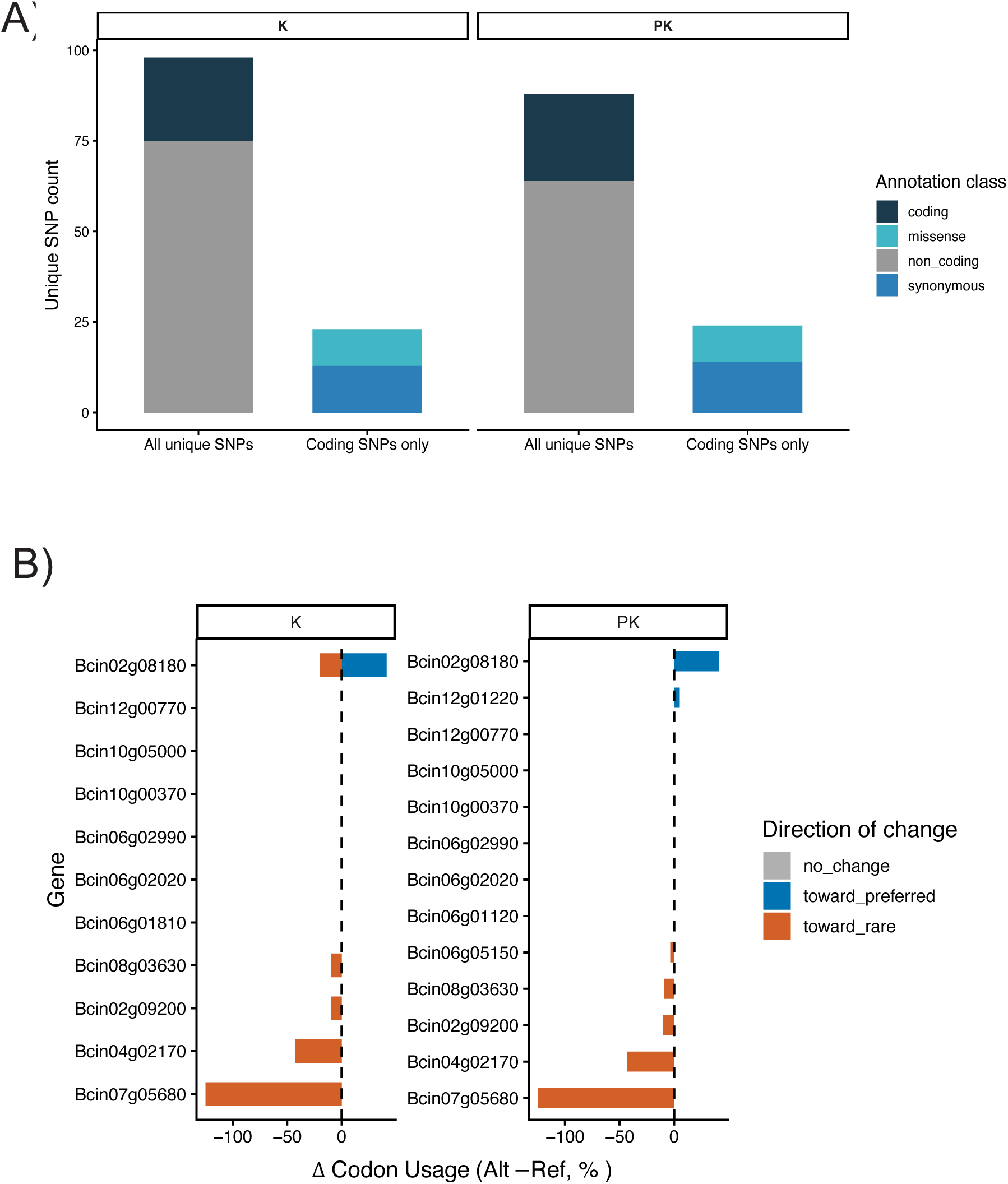
Functional annotation and codon bias shifts among genome-wide significant SNPs. (A) Distribution of genome-wide associated SNPs partitioned into coding and non-coding Categories in both the Kinship (K) model and Population Structure + Kinship (P+K) model. Coding SNPs are further subdivided into synonymous or missense variants. (B) Change in codon usage bias for synonymous SNPs in coding regions between the alternate and reference alleles. Positive values indicate shifts toward more preferred codons, whereas negative values indicate shifts towards non-preferred codons. Colors denote the direction of change, and gene labels indicate the trait class.

Among synonymous SNPs, codon usage frequency varied in direction and magnitude (Figure 4B). 15 variants (60 %) shifted towards rarer codons, three variants (12%) shifted towards the preferred codon, and seven variants (28%) shifted to codons with approximately equal usage across the genome. At the gene level, synonymous hits shifted toward rarer codons (4/11), suggesting a potential directional bias in codon usage changes across multiple loci, with the largest magnitude change observed in *Bcin07g05680* (MFS transporter encoding gene). In contrast, *Bcin02g08180* (AP complex subunit beta) exhibited a synonymous shift in which the minor allele encoded a more preferred codon relative to the major allele (Figure 4B).

### Docking Results

Bcin07g05680 was selected as a protein of interest for follow-up docking analysis, based on the presence of significantly associated synonymous SNPs. This gene was selected as a candidate because its functionality is expected to be conserved across the population, while its expression may be more sensitive to transcriptional and/or translational variation arising from codon usage differences^57^. Putative ligand transport pathways were identified using CaverWeb to calculate potential access tunnels. CaverWeb generated over 232 pockets of interest. As many of these predicted pathways were either shallow, discontinuous, or sterically constrained, subsequent analyses focused on tunnels likely to represent physiologically relevant transport routes. The primary pocket was chosen based on quantitative criteria associated with functional substrate channels, such as having the widest bottleneck (2.87 ± 0.43), the shortest path (11.46 ± 5.79), and a curvature of 1.14 (± 0.11). These parameters suggested a more accessible path for ligand transport than other tunnels.^41^ Using Caverweb, linalool was docked to transport through the first tunnel. Transport through this tunnel exhibited a favorable trajectory, with values ranging from approximately -4.0 kcal/mol near the internal binding region to -2.6 kcal/mol as it exited the pocket. This trajectory suggests a low energetic barrier for diffusion through the tunnel, supporting the possibility that this transporter could facilitate the export of small, lipophilic compounds such as linalool.

## Discussion

### Phenotypic Variation

Filamentous fungi exhibit substantial phenotypic plasticity in growth form and metabolism in response to environmental stressors, and these responses often reflect alternative strategies that balance exploration, resource acquisition, and damage mitigation^2,12^. In *B. cinerea,* exposure to linalool revealed isolate-to-isolate variation in metabolic activity, as quantified by ΔE and b* dynamics (Figure 1B). Isolates with steep ΔE slopes displayed rapid color shifts, suggesting rapid metabolic activity. Isolates with gradual slopes may experience a slower metabolic response or higher susceptibility to linalool toxicity. Linalool has been shown to disrupt mitochondrial function and downregulate mitochondrial and ergosterol genes, as well as glutathione synthesis^65^, which may underlie the observed differences in ΔE trajectories.

These patterns were recapitulated by PCA of metabolic traits (Figure 1B). PC1 (63.8% of total variance) captured response magnitude, separating isolates with higher AUC and steep ΔE slopes (fast responders) from those with more gradual slopes (slow responders). PC2 (16.7%) captured variation in response timing and trajectory, distinguishing isolates with earlier versus later dye reduction. Although isolates formed two broad groups with minimal overlap along PC1, overlap among PC2 suggests that metabolic phenotypes are better described as a continuum dominated by response strength rather than discrete classes. Together, these patterns suggest that isolates differ in both the strength and the rate at which they maintain metabolic reducing capacity under linalool exposure, potentially reflecting variation in tolerance mechanisms. Similarly, Fusarium responds to linalool exposure through broad transcriptional remodeling, with enrichment in ABC transporter activity, sterol biosynthesis, and glutathione metabolism^37^, supporting a model in which linalool perturbs membrane homeostasis, detoxification, and redox buffering pathways. While resistance to chemical stress can be expressed not only through physiological detoxification. Filamentous fungi can also respond by reprogramming growth through adjustments in hyphal tips, branching, and network connectivity to reallocate growth under inhibitory conditions. In *Rhizophagus irregularis,* exposure to fungicides reduces hyphal length and alters anastomosis, demonstrating that chemical environments can reshape hyphal architecture and interaction behavior^49^.

Isolates not only differ in metabolic activity but also in structural traits, reflecting differences in how each isolate’s hyphal network responds to linalool (Figure 1A). Network features, such as fractal dimension, hyphal tips, and edges, varied across the population, reflecting differences in network complexity and connectivity (Supplementary Table 1). These traits reveal that some isolates form compact, highly interconnected networks, whereas others exhibit more of an exploratory phenotype. In the structural PCA (Figure 1A), isolates are separated into three broad morphological groupings driven by network size and connectivity metrics (edges, nodes, tips, branches, and total hyphal length). Similar contrasts have been described in filamentous fungi under different environmental conditions, where dense, cohesive networks are associated with nutrient-rich environments and exploratory growth with nutrient poor environments^2,23^. In the context of linalool, one possibility is that highly connected networks increase physiological robustness (i.e., maintaining growth by distributing transport and reducing power demands across a denser mycelial system), while exploratory architectures reflect avoidance or slower growth under stronger inhibition. More broadly, the observed architectural plasticity aligns with evidence that *B. cinerea* can undergo hyphal specialization as a strategy to overcome host defenses^48^. Infection cushions, specialized multicellular structures that secrete cell wall-degrading enzymes, exhibit higher tolerance to host defenses^48^. The structural diversity we observe may mirror this, suggesting that remodeling of the hyphal network is a key in *B. cinerea’s* ability to overcome host defenses and enact disease.

### Genetic Architecture of Resistance to Linalool

Linalool exposure is known to elicit broad physiological remodeling in *B. cinerea,* including disruption of mitochondrial function and changes in sterol and redox processes^1,65^. In a proteomic time course, brief exposure to linalool led to broad proteomic changes, involving enrichment of proteins involved in translation, amino acid metabolism, and post-translational modification^65^. These findings suggest that linalool triggers energetic and metabolic reprogramming in response to mitochondrial stress and membrane disruption.

In our GWAS, the strongest metabolic response associations localized to *Bcin07g05680*, an MFS transporter-encoding gene. Most of these variants were synonymous, indicating no change in the amino acid sequence; however, this mutation could influence protein expression or conformation via mRNA stability or translation efficiency^34^. This mechanism has been demonstrated in the multidrug resistance gene MDR1, where a synonymous polymorphism altered substrate specificity by altering folding and drug-protein interactions^34^. The synonymous mutations in *Bcin07g05680* resulted in shifts toward the use of non-preferred or uncommon codons (Figure 4B). Importantly, codon usage mismatch has been shown to have strong functional consequences in *B. cinerea*^57^. Codon optimization of the geneticin resistance gene NPTII to match *B. cinerea* codon preferences increases transformation by 30-fold, demonstrating that *B. cinerea* is sensitive to codon usage in heterologous coding sequences^57^. While a shift toward rarer codons could be detrimental under baseline conditions due to reduced translational efficiency, the net fitness effect of these synonymous changes may be context-dependent and could be favored under linalool exposure^44^. In *Salmonella,* codon usage bias has been linked to stress and virulence-associated gene regulation, where enrichment towards synonymous codons is thought to support translation under nutrient-limited conditions and allows for colonization of different host tissues^11^.

Efflux transporters are frequently implicated in chemical stress responses in fungal pathogens. In Fusarium, a GWAS examining sensitivity to the demethylation inhibitor (DMI) class of fungicides identified candidate loci mapping to transport-related proteins^56^, suggesting that membrane transport is part of a broader genetic toolkit to cope with chemical-related stress. In *B. cinerea,* for example, transcriptomic comparisons of fludioxonil-resistant isolates found MFS to be upregulated in fludioxonil-resistant isolates^1^. In addition, the *MfsG* transporter has been identified as a virulence factor that confers tolerance to other SMs, such as glucosinolates^61^. To explore whether this transporter could interact with linalool, molecular docking was performed using the predicted *Bcin07g05680* (MFS transporter) protein structure. Notably, the variants in/near *Bcin07g05680* (MFS transporter) *were* predominantly synonymous and shifted codon usage toward rarer codons, providing a plausible translation-linked route by which silent variation could influence transporter performance. In addition, *Bcin07g05210,* a btb domain containing protein, was associated with variation in metabolic response, suggesting a potential regulatory role in mediating cellular adaptation to linalool exposure.

Several SNPs were linked to variation in structural traits, including the area under the fractal dimension curve, branch fraction, and tip fraction (Figure 2). These traits capture differences in hyphal network connectivity and cohesion. Associated loci were in a mitochondrial translation initiation factor (*Bcin02g09200*), a Zn-C6 domain containing protein (*Bcin02g05460*), *Bcin02g08180* (AP complex subunit beta), and *bcltf1*. *Bcin02g09200* may contribute to variation in hyphal structure by regulating mitochondrial efficiency and availability. As hyphae extend and grow, the hyphal tips rely on ATP for growth; therefore, small shifts in mitochondrial activity could alter branching frequency and overall network cohesion. The ZN(II) C6 domain-containing transcription factor (Bcin02g05460) is likely to regulate downstream genes involved in carbon metabolism, stress signaling, and cell wall synthesis. In other filamentous fungi, ZN(II) C6 proteins regulate carbon metabolism and drug sensitivity^40^. *Bcin02g08180* encodes an AP complex subunit beta, which is involved in vesicle-mediated trafficking and cargo sorting. Vesicle trafficking is critical for fungal virulence in *B. cinerea* as it transports sRNAs to plants via extracellular vesicles^30^. Extracellular vesicles carry a diverse set of cargo that range from involvement in transport to autophagy^20^. *Bcltf1* has been categorized as a transcription factor that regulates virulence, with deletion mutants exhibiting hypersensitivity to oxidative stresses, therefore variation within *Bcltf1* could plausibly influence responses to oxidative stress from linalool exposure^52^.

Our *B. cinerea* population showed that in response to linalool exposure, phenotypic diversity is driven by metabolic and morphological plasticity. At the population level, metabolic resistance to linalool appears to be governed by a small number of genes, suggesting an oligogenic basis for this trait. This oligogenic pattern contrasts with prior studies of *B. cinerea* virulence and host, which have consistently shown highly polygenic architecutres^5,14^. Lesion formation and host specificity are influenced by hundreds of loci distributed through the genome, with many loci contributing small effects to overall pathogenicity ^5,14^. Polygenic architectures are expected for complex ecological traits that integrate host recognition, specialized metabolite production, nutrient acquisition, immune suppression, and growth across diverse host environments. In contrast, resistance to a specific host-derived defense compound, such as linalool, may depend more heavily on a limited number of genes with large functional effects. Candidate mechanisms may include transporters with broad substrate specificity, detoxification pathways, and general stress-response regulators that can rapidly alter cellular tolerance to chemical stress.

Consistent with our identification of an oligogenic architecture, previous experimental evolution studies have shown that exposure to specialized metabolites can select for multidrug resistance in *B. cinerea* via upregulation of ABC and MFS transporter genes^64^. Similarly, work in a Michigan *B. cinerea* population was associated with detectable population differentiation^46^. However, this study inferred population structure using partial sequences of a partial ITS sequences and a small number of loci (*G3DPH*, *HSP60* and *RPB2),* rather than genome-wide variation. While these sequences are useful for species identification and broad phylogenetic placement, they do not capture genome-wide variation and offeroffer relatively low resolution for detecting population structure within highly recombining populations such as *B. cinerea*^7,14^.

In contrast, our study leveraged whole-genome sequencing data, allowing substantially finer-scale inference of genetic relatedness and population structure. Despite isolates originating from diverse geographic regions and host species, we observed remarkably little population subdivision across the panel (Supplementary Figure 2). This finding supports previous evidence that *B. cinerea* functions as a broadly recombining generalist pathogen with limited host-level genomic specialization. The lack of strong host- or geography-associated structure further suggests that adaptive responses to specialized metabolites such as linalool are unlikely to be restricted to isolated evolutionary lineages. Instead, resistance-associated alleles may spread broadly across interconnected populations or repeatedly arise under chemical selection. Together, our results suggest that while virulence in *B. cinerea* is governed by a highly polygenic architecture, adaptation to specific host-derived chemical defenses may occur through selection on a comparatively small number of loci with large phenotypic effects.

Across our study, results provide insight into how *B. cinerea* overcomes host chemical defenses and how resistance to specialized metabolites integrates with broader virulence mechanisms. Our findings suggest that the success of *B. cinerea* as a generalist pathogen may arise from a mixed genetic architecture in which highly polygenic virulence traits are coupled with comparatively oligogenic resistance mechanisms to specific host-derived compounds, such as linalool. This genetic architecture may allow *B. cinerea* to maintain broad virulence across hosts while also rapidly adapting to localized chemical environments through selection on a relatively small number of high-effect resistance loci. Although this study represents one of the first population-level investigations of *B. cinerea* responses to linalool, our isolate panel captures only a subset of the species’ global genetic diversity and therefore may not encompass the full allelic diversity underlying resistance phenotypes. In addition, isolates were exposed to a single VOC, whereas natural host-pathogen interactions involve complex and dynamic mixtures of hundreds of specialized metabolites that vary across host species, tissues, developmental stages, and infection conditions. Consequently, the resistance mechanisms identified here likely represent only one component of the broader chemical interactions shaping pathogen success in natural systems. Future studies incorporating larger, more globally representative collections of isolates, combined with exposure to chemically diverse metabolite mixtures, will be necessary to resolve the full genetic architecture underlying resistance to host-derived defenses. More broadly, integrating population genomics with metabolomics and experimental evolution approaches may help clarify how generalist pathogens maintain the capacity to tolerate chemically heterogeneous host environments while retaining high levels of virulence across diverse plant species.

## Data Availability Statement

All genomic data used in this study was generated by Caseys et al^14^.All code used for image processing and downstream analyses is available at (https://github.com/melaniemadrigal07/GWASLIN). Supplementary material is available at 10.5281/zenodo.19425171 and provides detailed information supporting the analyses performed in this study. Supplementary Table 1 provides descriptions of the traits used in this study. Supplementary Table 2 contains trait means. Supplementary Table 3 contains results of the genome wide association analyses it includes SNPs, genomic positions, effect sizes, and statistically significant values for all trait-SNP associations. Supplementary Table 4 provides descriptions of all annotation fields used in SNP characterization including genomic location, allele identity, codon context, and functional classification. Supplementary Table 5 contains location of the SNP and classification of the mutation.

## Acknowledgments

J.A.D., M.M., and J.C.M. were funded in part by a USDA grant (2022-67012-43019). M.M. was also funded by the Future Scholar fellowship at Louisiana State University.

M.M. and J.A.D. designed the study; M.M. and J.C.M. collected the data; M.M, J.C.M., and J.A.D. analyzed the data; and M.M. and J.A.D. wrote the manuscript with input from D.J.K. and J.C.M.

