## Supplemental Figures for "Strategy Sets the Scene: Genetic architecture of linalool resistance in *Botrytis cinerea*"

Figure S1. Pairwise relationships among all measured traits shown as scatterplots, density distributions, and Pearson correlation coefficients. The lower triangle contains scatterplots illustrating pairwise relationships between traits. The diagonal panels display density plots of each individual trait. The upper triangle shows Pearson correlation coefficients for each trait pair, providing the strength and direction of linear associations among traits.
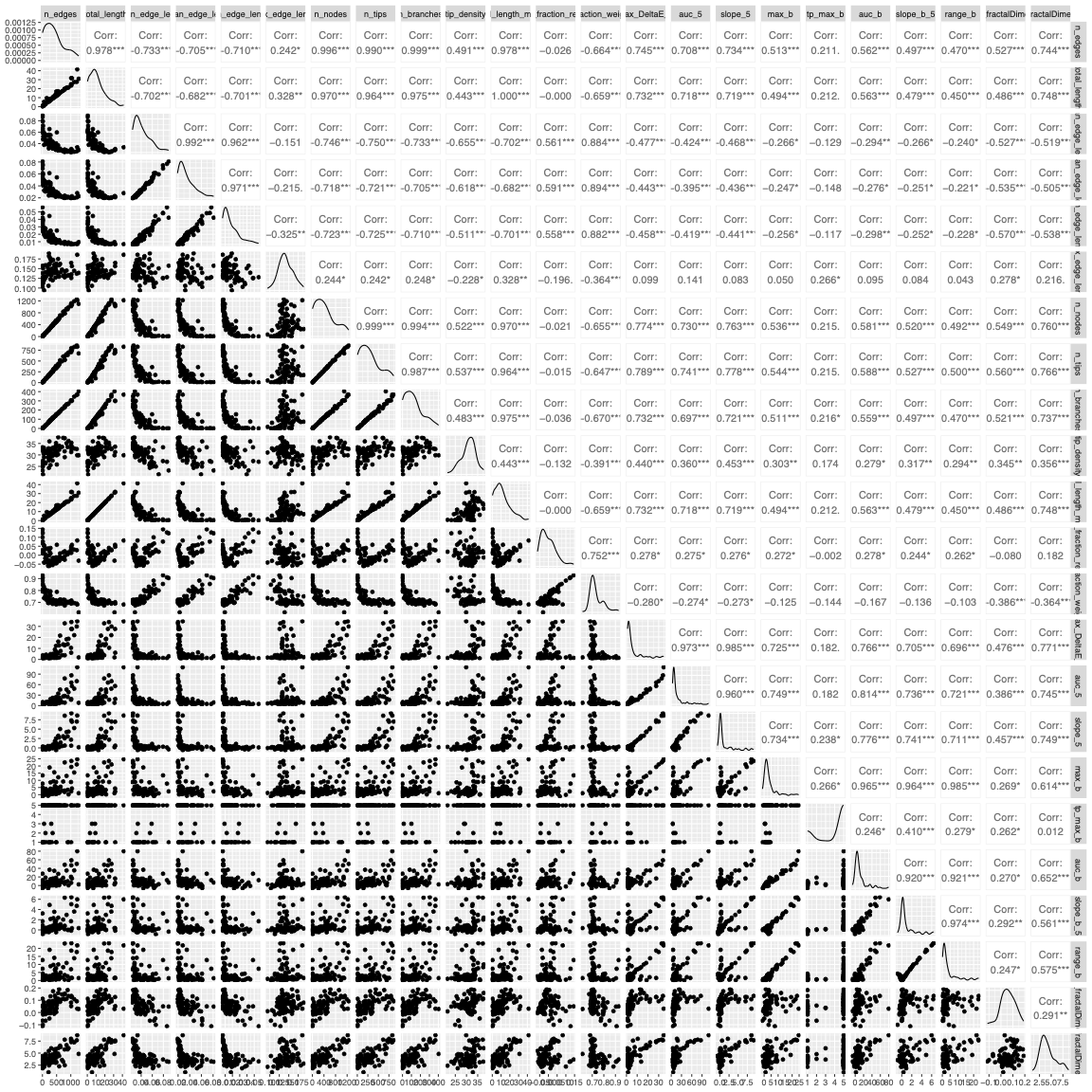

Figure S2. Principal Component Analysis of *Botrytis cinerea* population(A-C). Isolates are colored according to host of origin and shaped according to geographic region. (D) Scree plot used to identify how many PC axis to retain to capture population structure

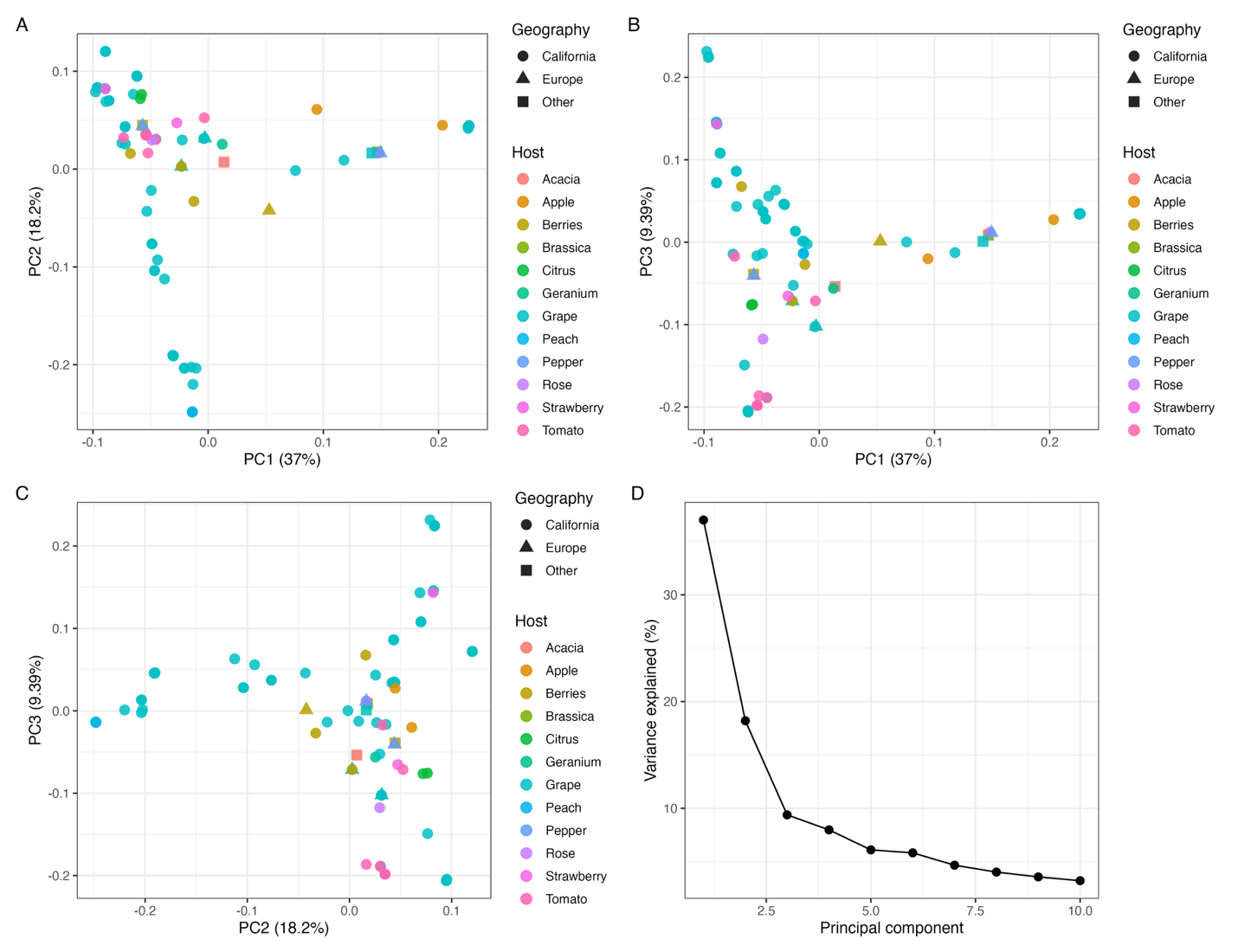

Figure S3. Quantile-Quantile plots of genome wide association mapping results for metabolic and structural traits: (a) area under the curve of ∆E at timepoint 5, (b) area under the curve of b* at timepoint 5, (c) max b*, (d) max ∆E, (e) slope of ∆E at timepoint 5, (f) slope of b* at timepoint 5, (g) range of b*, (h) maximum edge length, (i) minimum edge length, (j) median edge length, (k) area under the curve of fractal dimension, (l) slope of fractal dimension, (m) number of hyphal tips (n) number of hyphal nodes, (O) tip fraction residuals,(p) weighted tip fractions,(q) weighted tip fractions,(r)mean total hyphal length,(s) tip density. The shaded region indicates the 95% confidence interval around the expected null distribution (red reference line). Observed values from the kinship-corrected model (K) are shown in orange, whereas values from models incorporating both kinship correction and population structure (P+K) are shown in green.

a) area under the curve of ∆E at timepoint 5 (20 hours post inoculation)

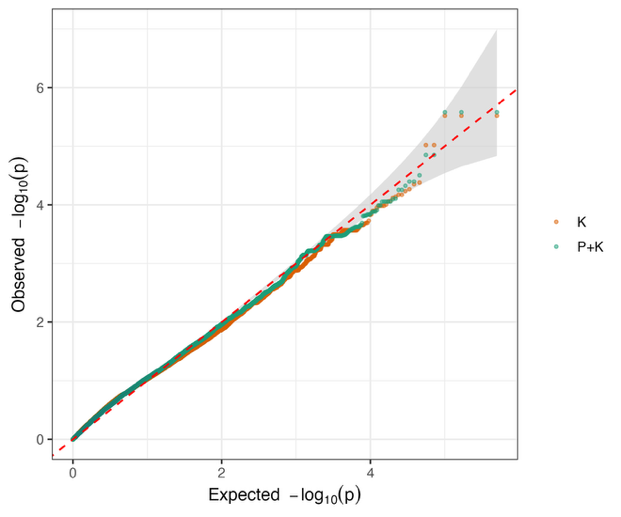

b) area under the curve of b* at timepoint 5 (20 hours post inoculation)

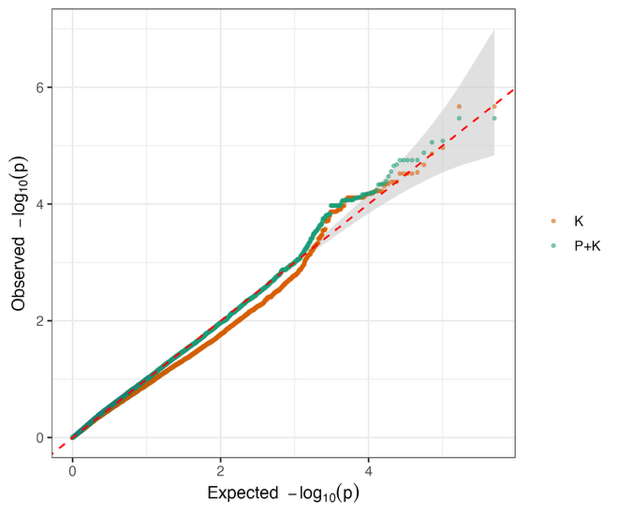

c) max b*

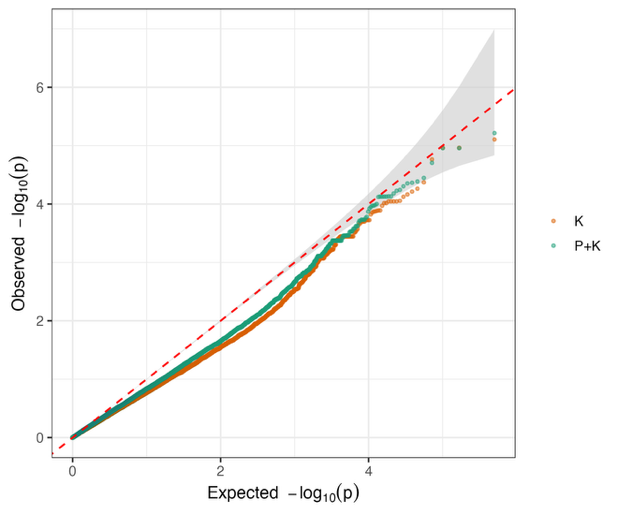

d) max ∆E

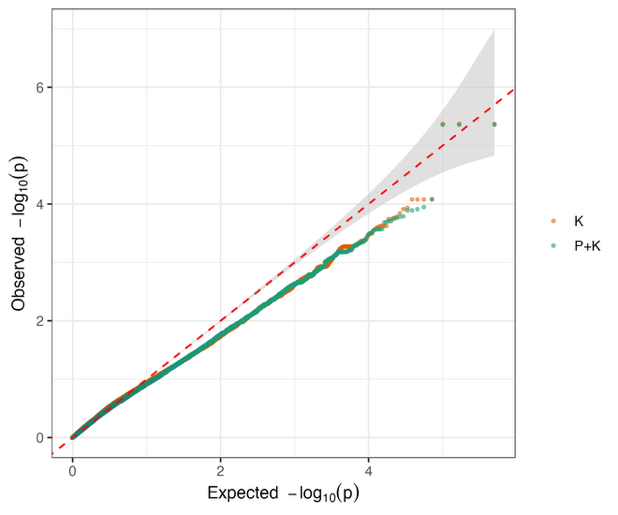

e) slope of ∆E

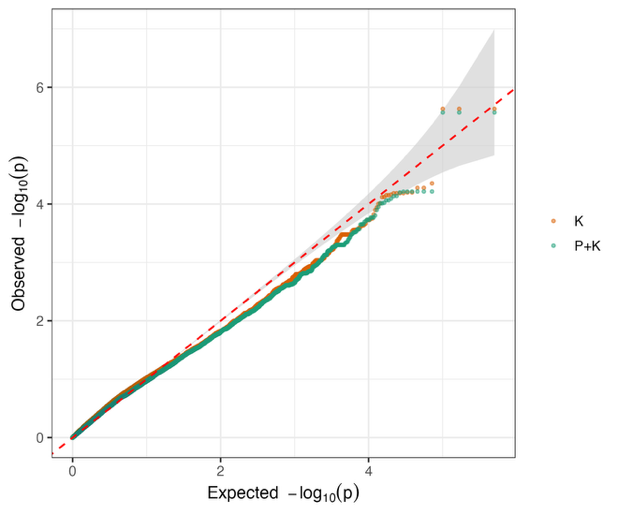

f) slope of b* at timepoint 5 (20 hours post inoculation)

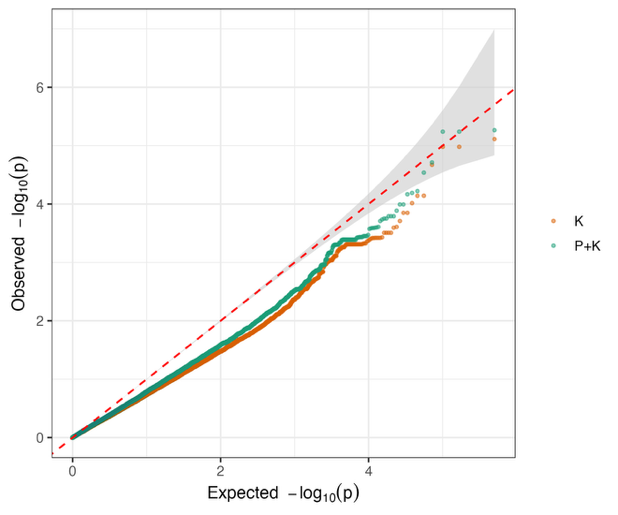

g) range of b*

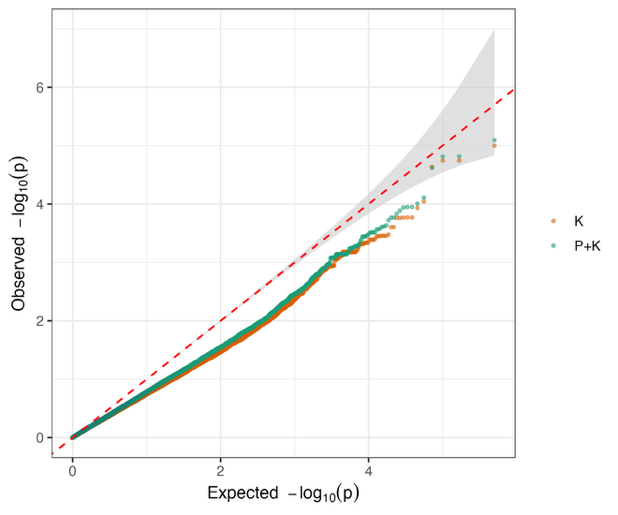

h) maximum edge length

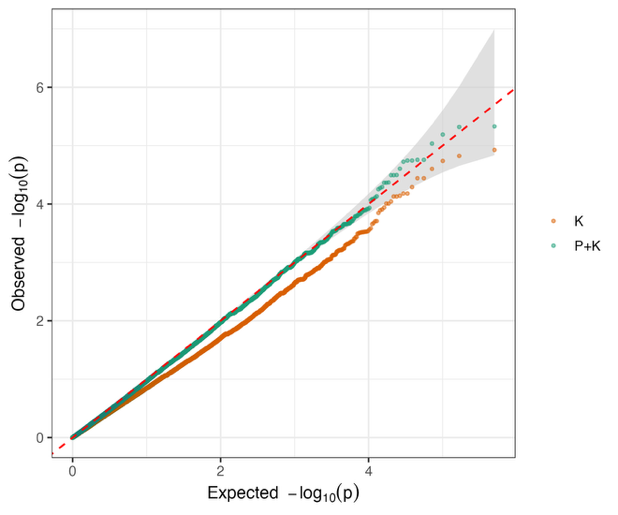

i) minimum edge length

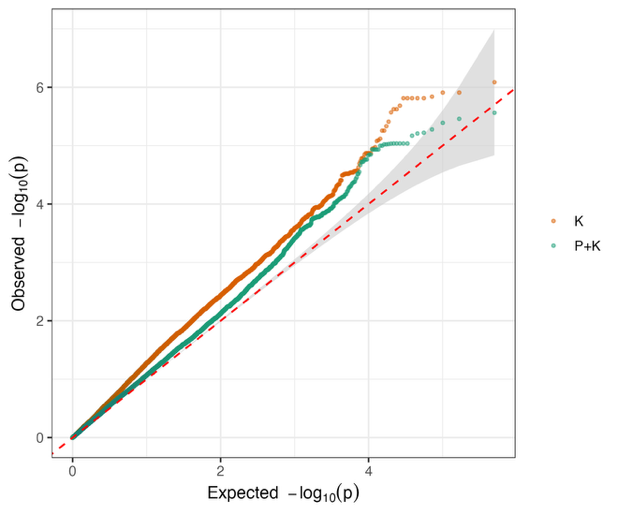

j) median edge length

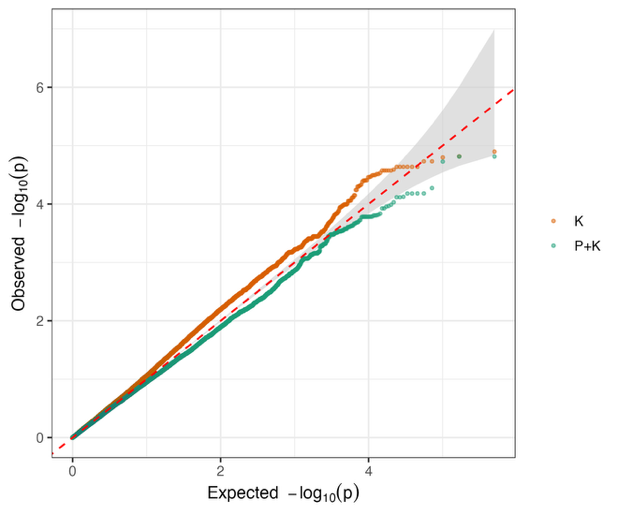

k) area under the curve of fractal dimension

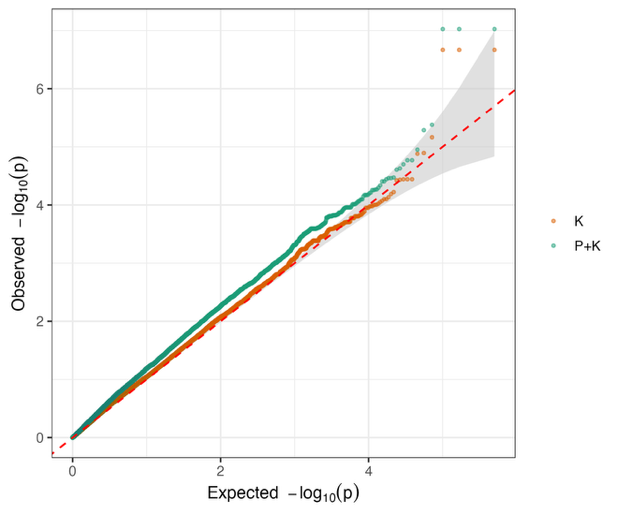

l) slope of fractal dimension

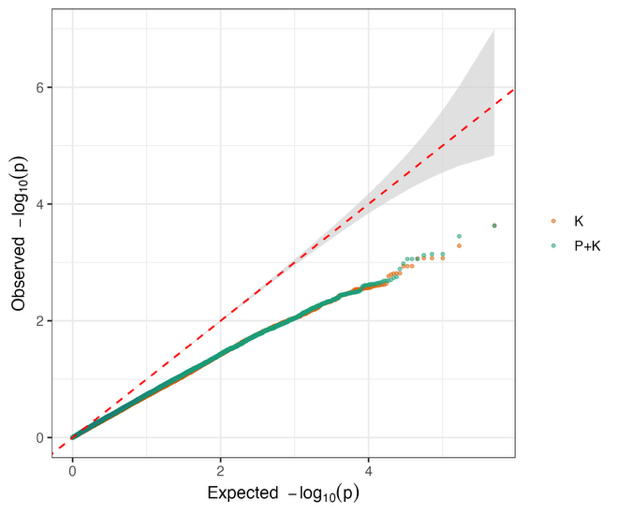

m) number of hyphal tips

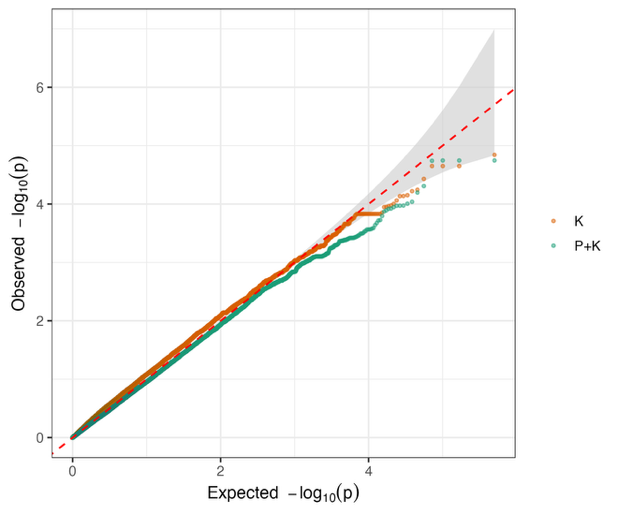

n) number of hyphal nodes

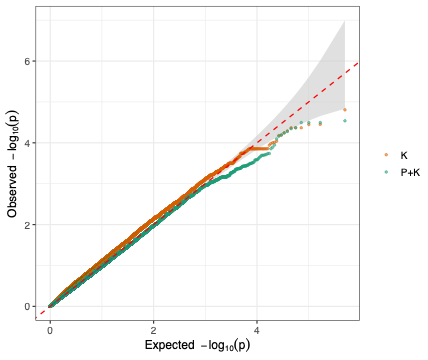

o) tip fraction residuals

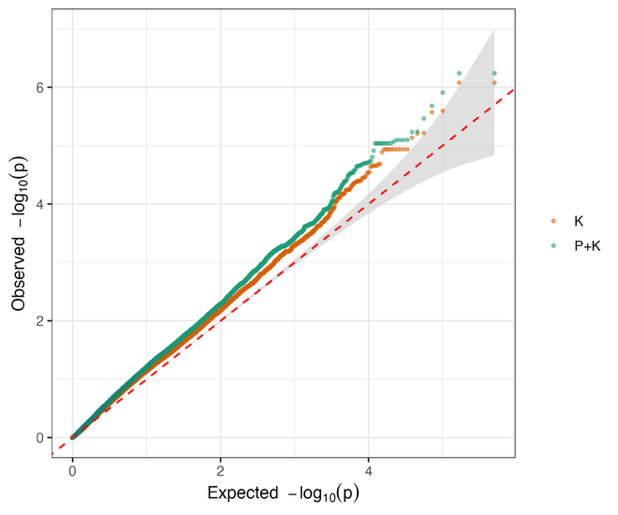

p) weighted tip fractions

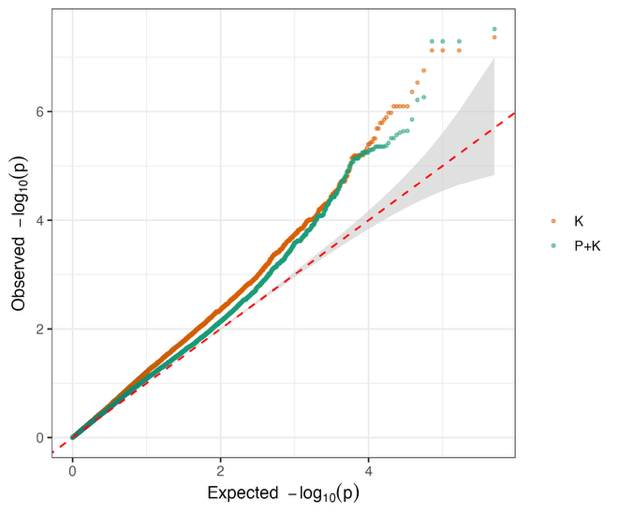

q) weighted tip fractions

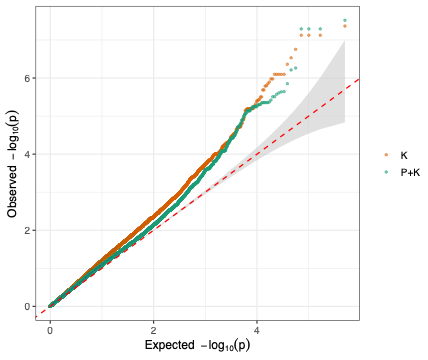

r) total hyphal length mean

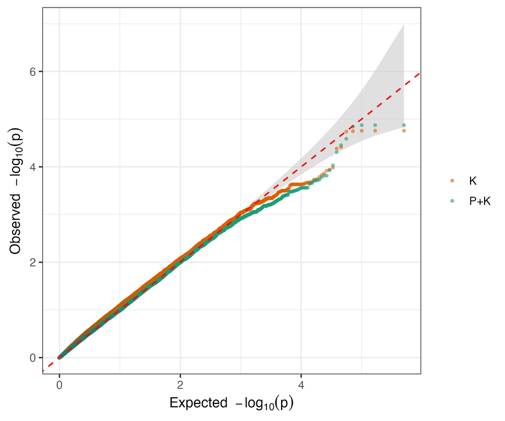

s) tip density

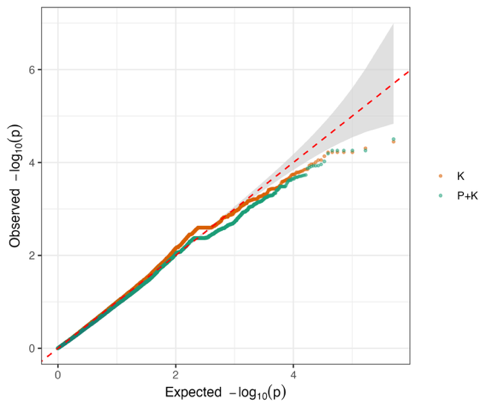

t) branch fraction

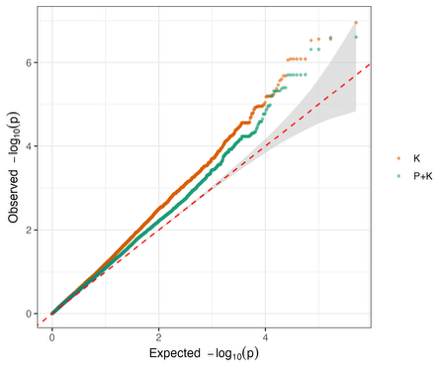

Figure S3. Manhattan plots of genome-wide association mapping for all traits: (a) area under the curve of ∆E at time point 5, (b) area under the curve of b* at timepoint 5, (c) max b*, (d) max ∆E, (e) slope of ∆E at timepoint 5, (f) slope of b* at timepoint 5, (g) range of b*, (h) maximum edge length, (i) minimum edge length, (j) median edge length, (k) area under the curve of fractal dimension, (l) slope of fractal dimension, (m) number of hyphal tips (n) number of hyphal nodes, (O)   tip fraction residuals, and (p) weighted tip fractions,(q) tip density ( r) total hyphal length, (s) branch fraction. The red threshold line refers to the Bonferroni significance threshold. Values from the kinship-corrected model (K) are shown in blue, whereas values from models incorporating both kinship correction and population structure (P+K) are shown in purple. Gene names for the nearest gene to SNPs above the Bonferroni threshold are labeled with the model color in which the association was identified.

(a) area under the curve of ∆E at time point 5

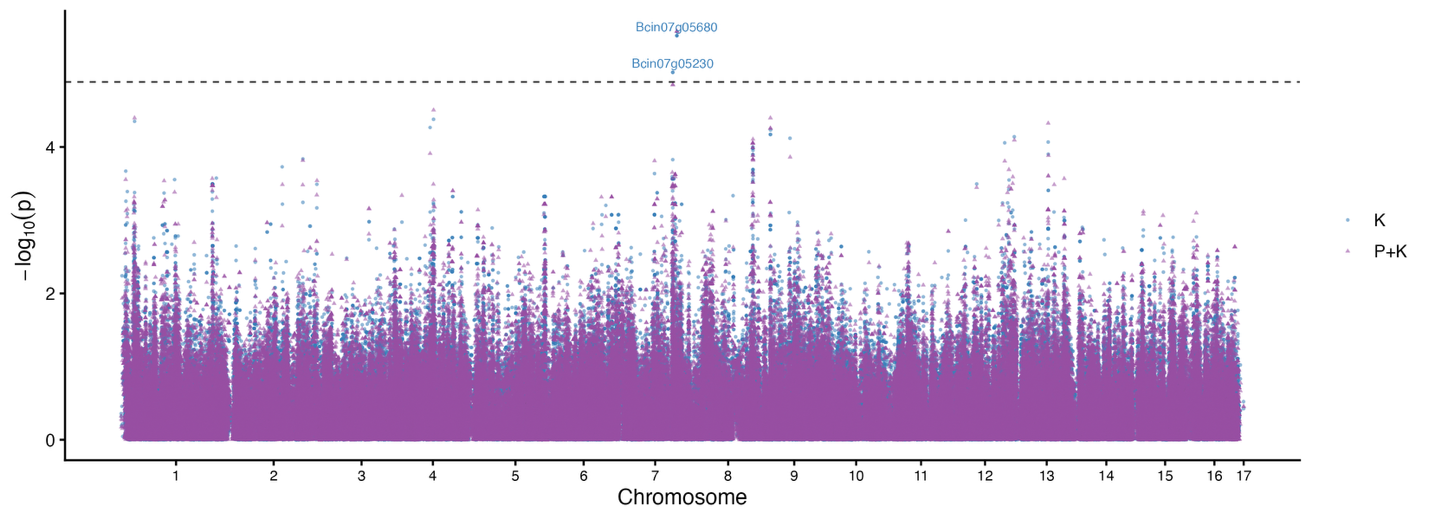

(b) area under the curve of b* at timepoint 5,

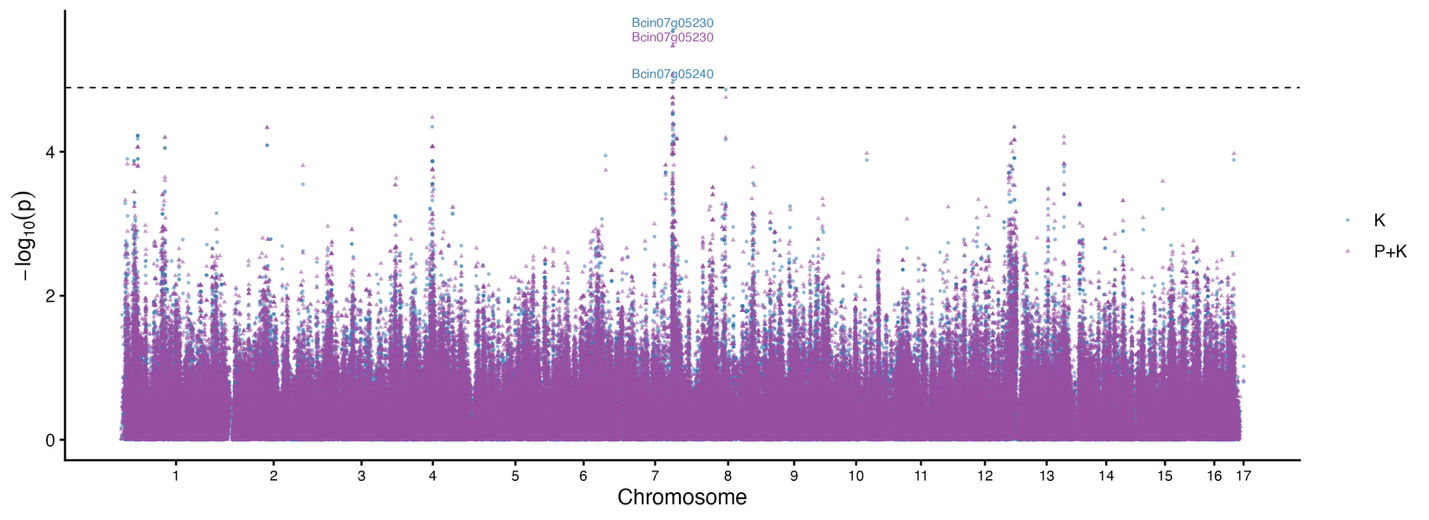

(c) max b*
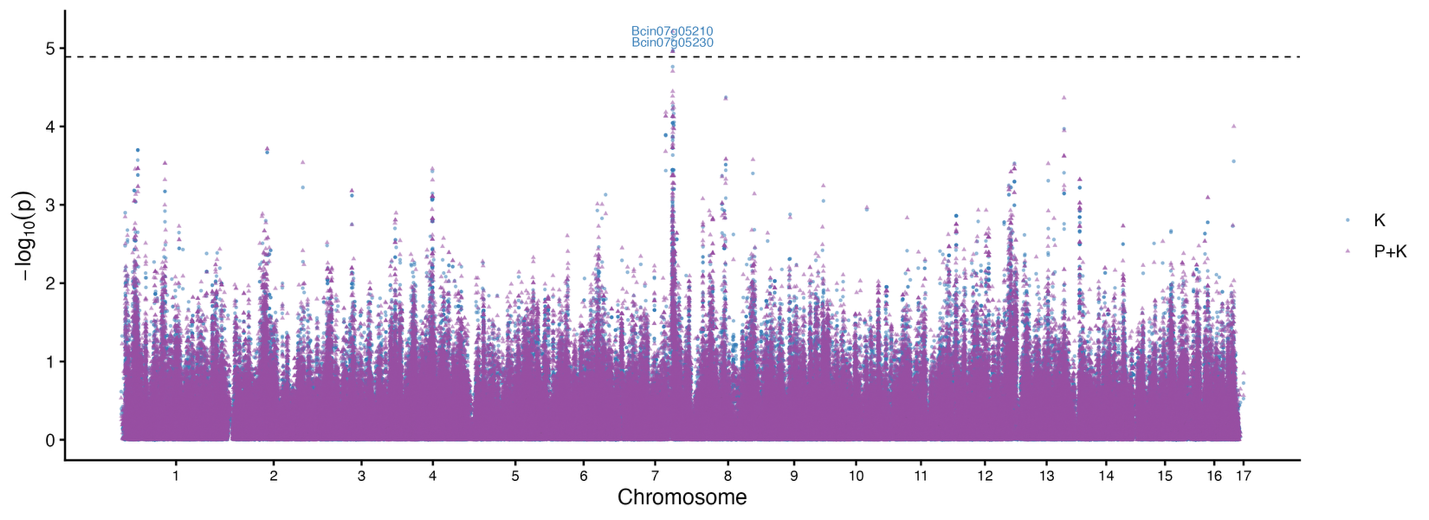

(d) max ∆E

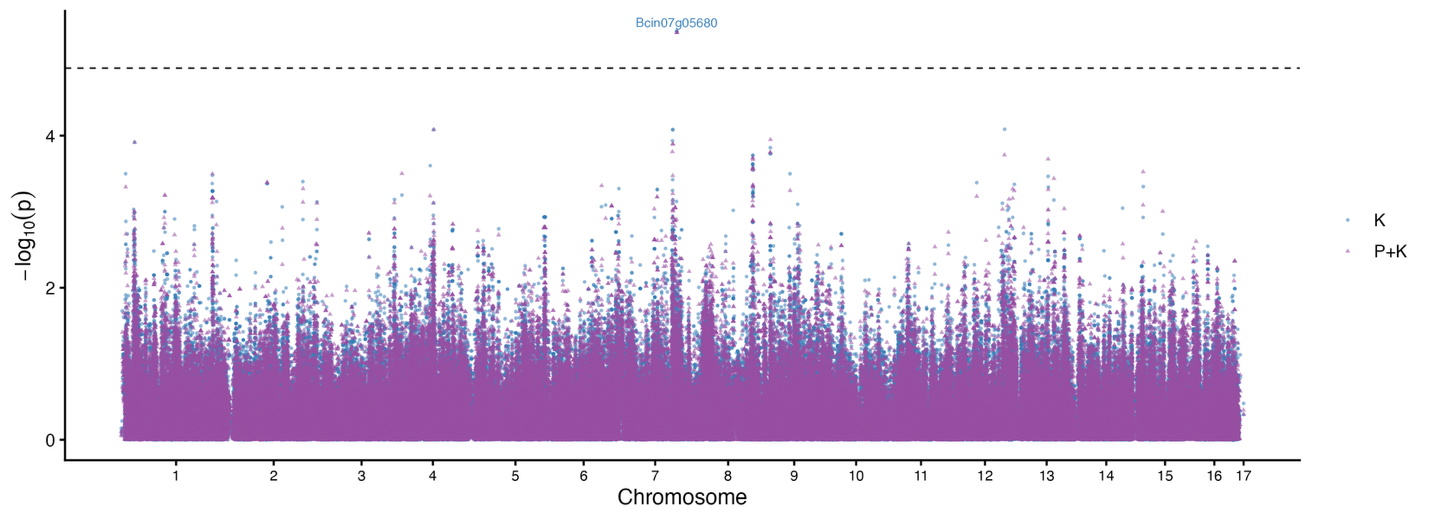

(e) slope of ∆E at timepoint 5

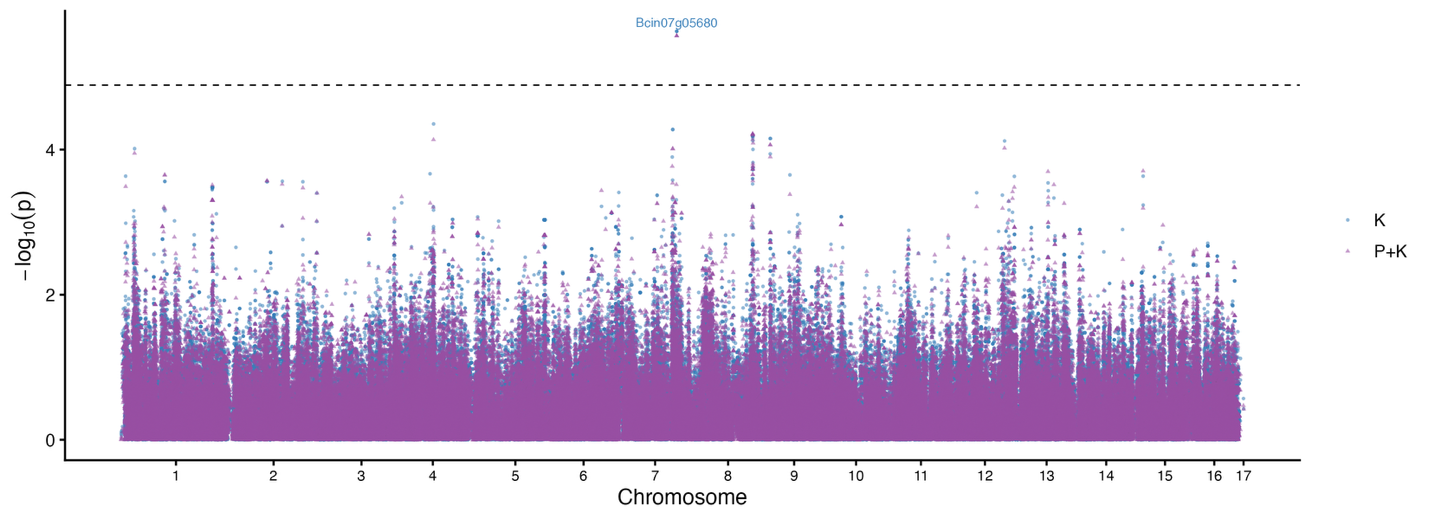

(f) slope of b* at timepoint 5

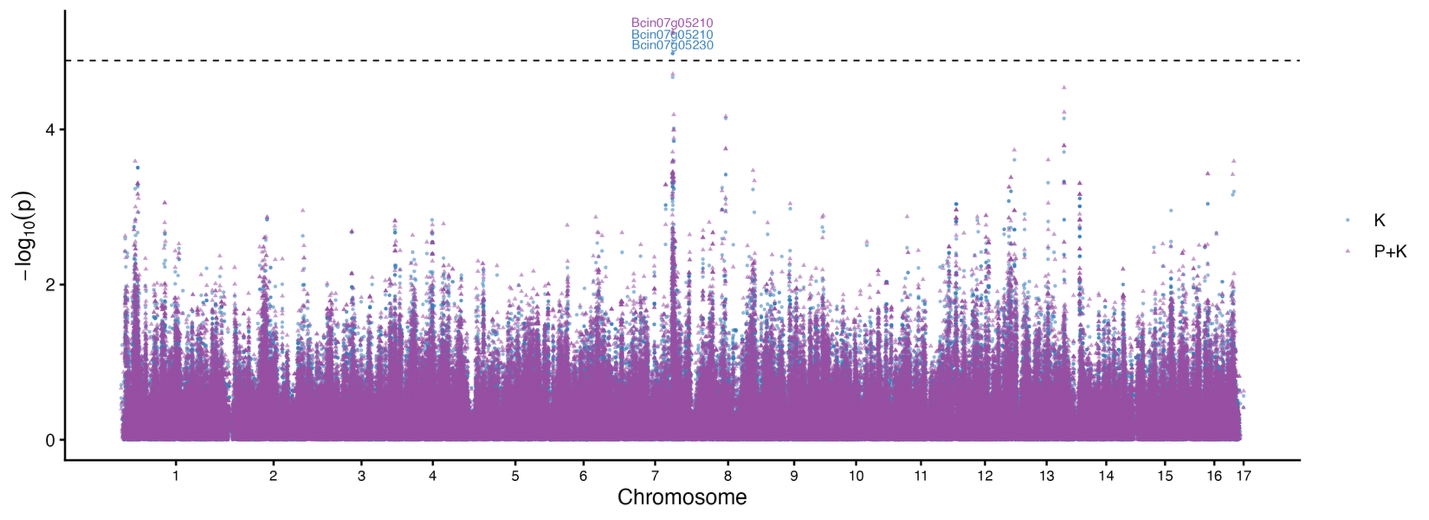

(g) range of b*

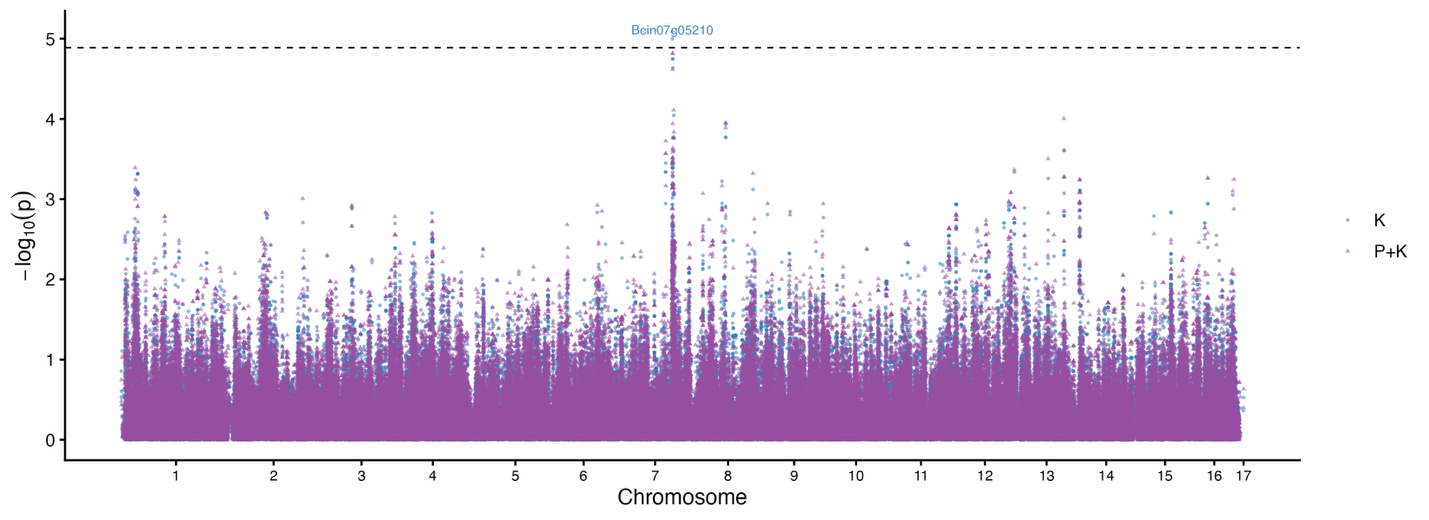

(h) maximum edge length

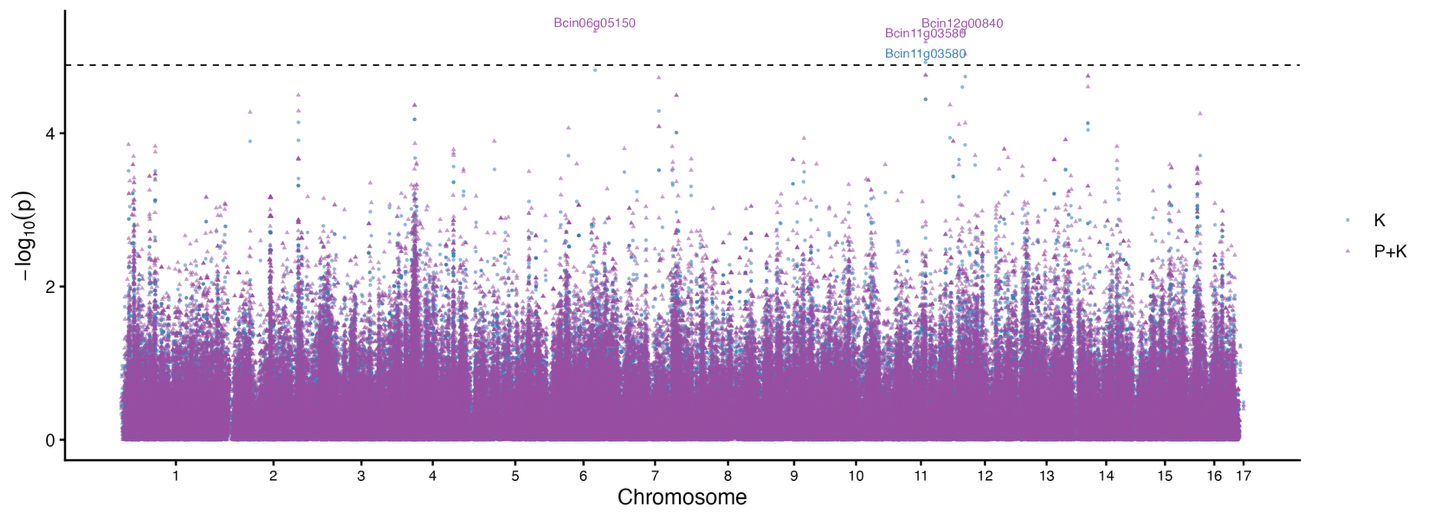

(i) minimum edge length

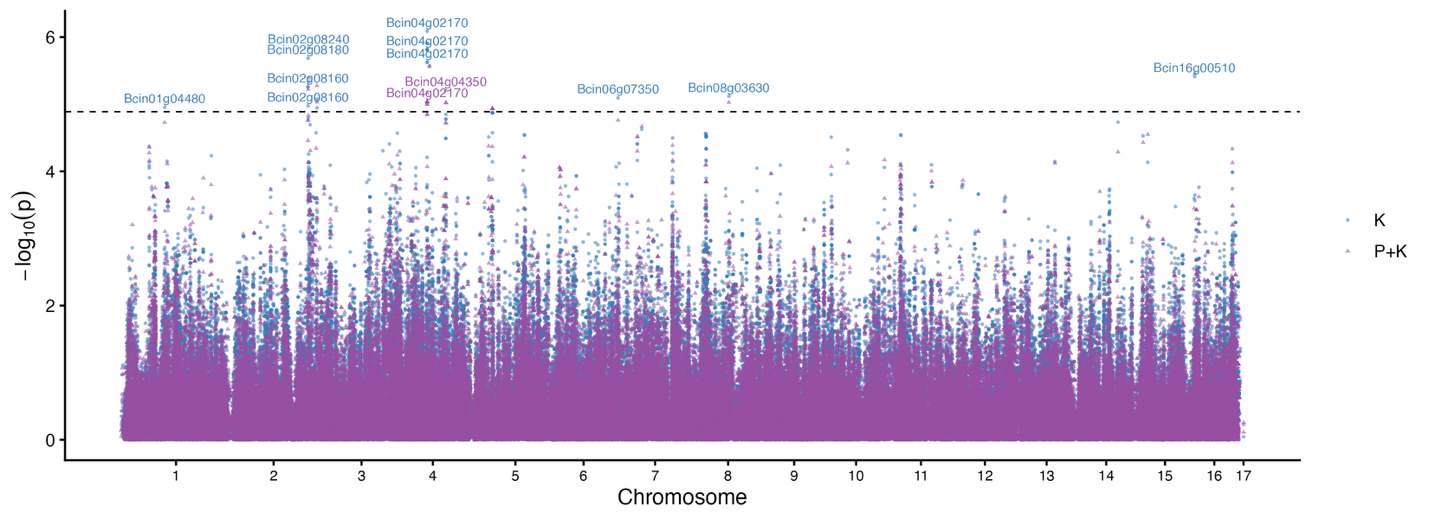

(j) median edge length

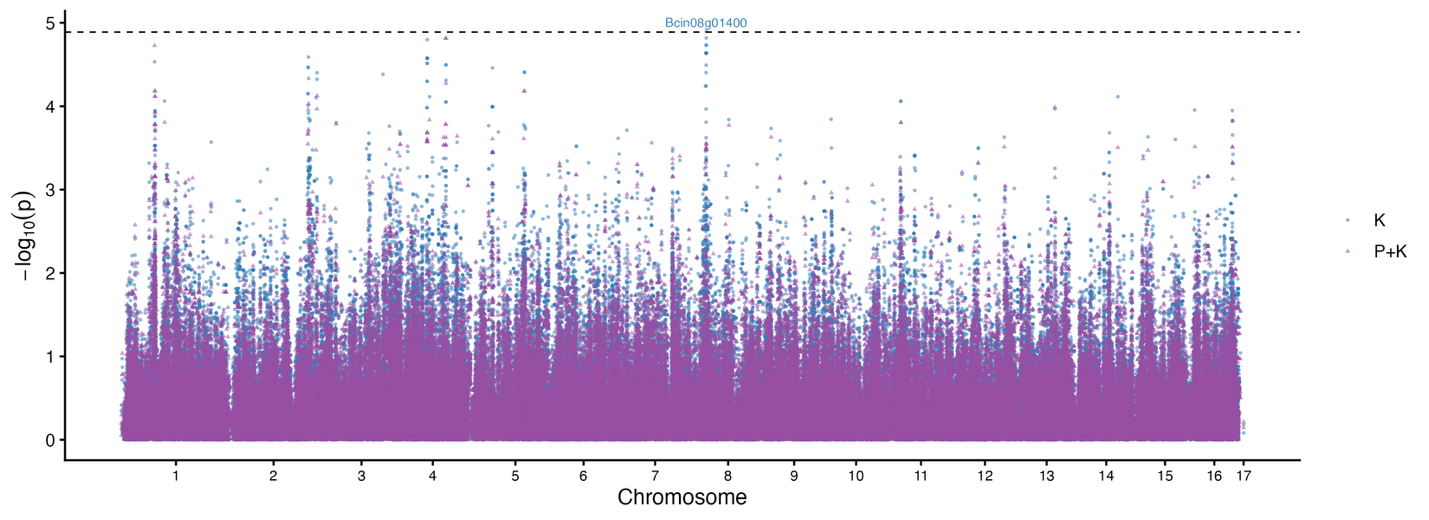

(k) area under the curve of fractal dimension

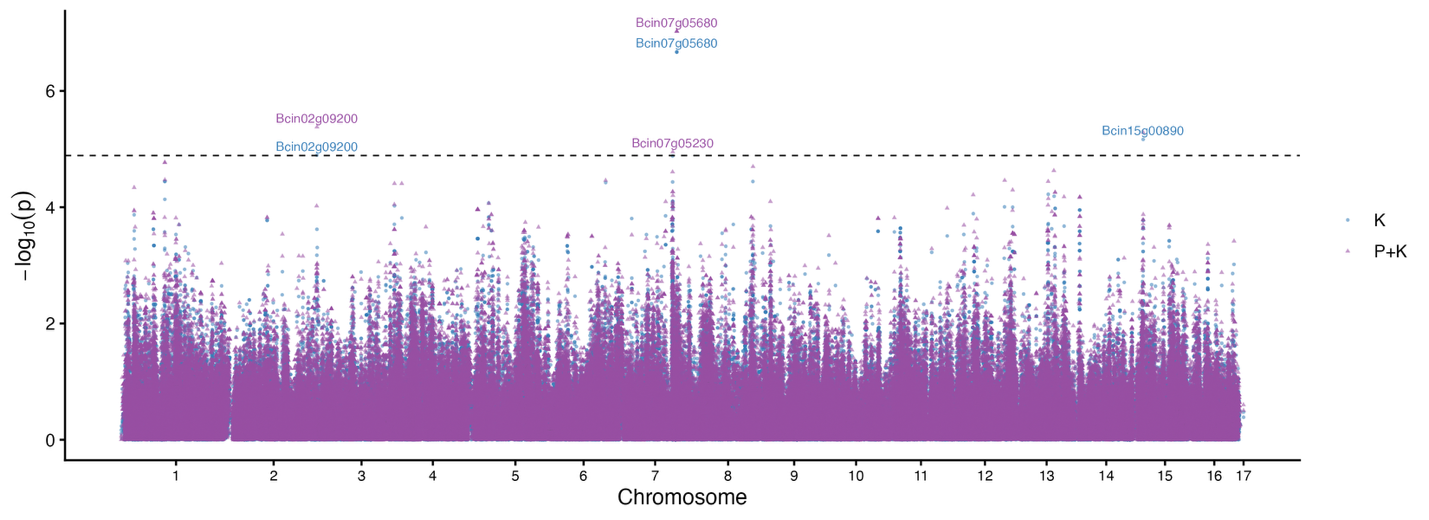

(l) slope of fractal dimension

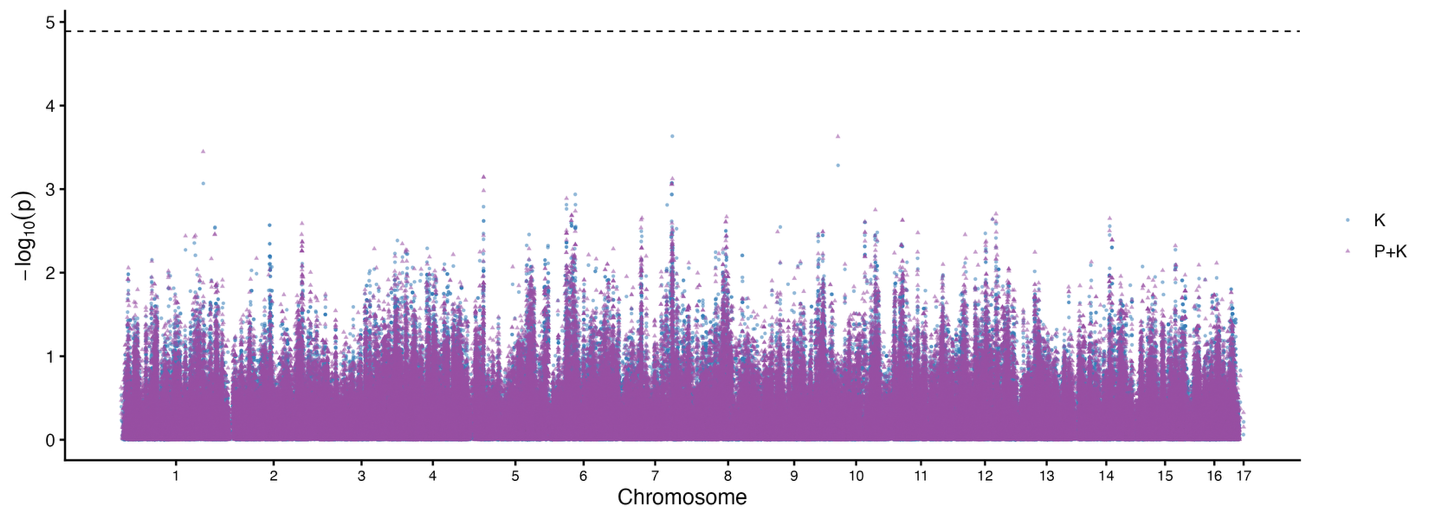

(m) number of hyphal tips

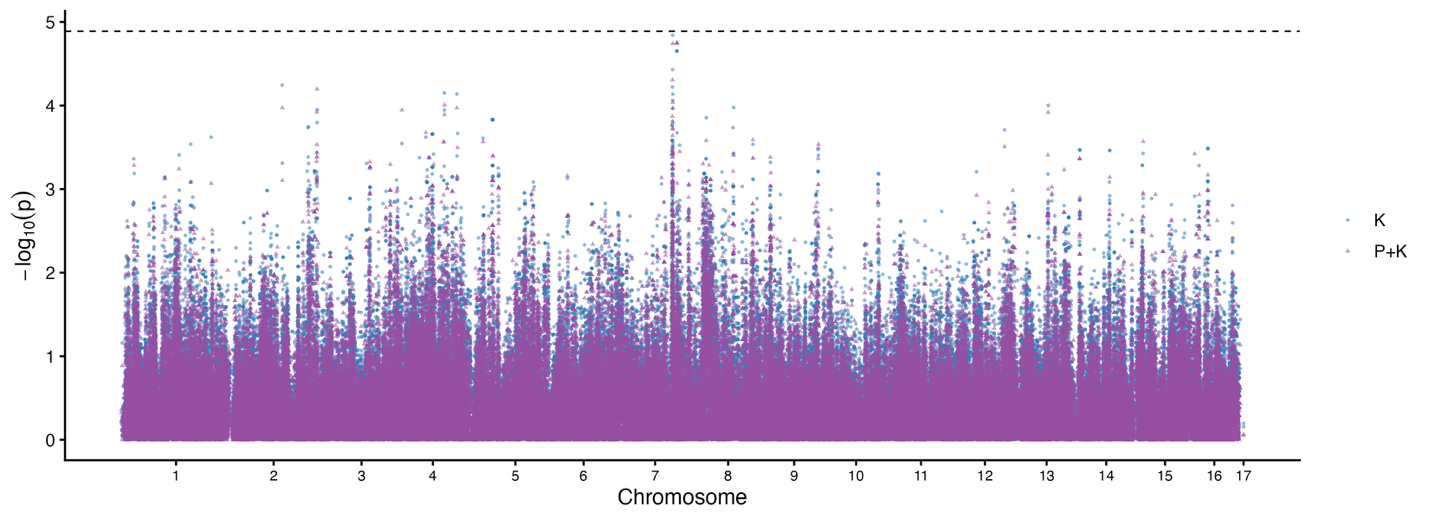

(n) number of hyphal nodes

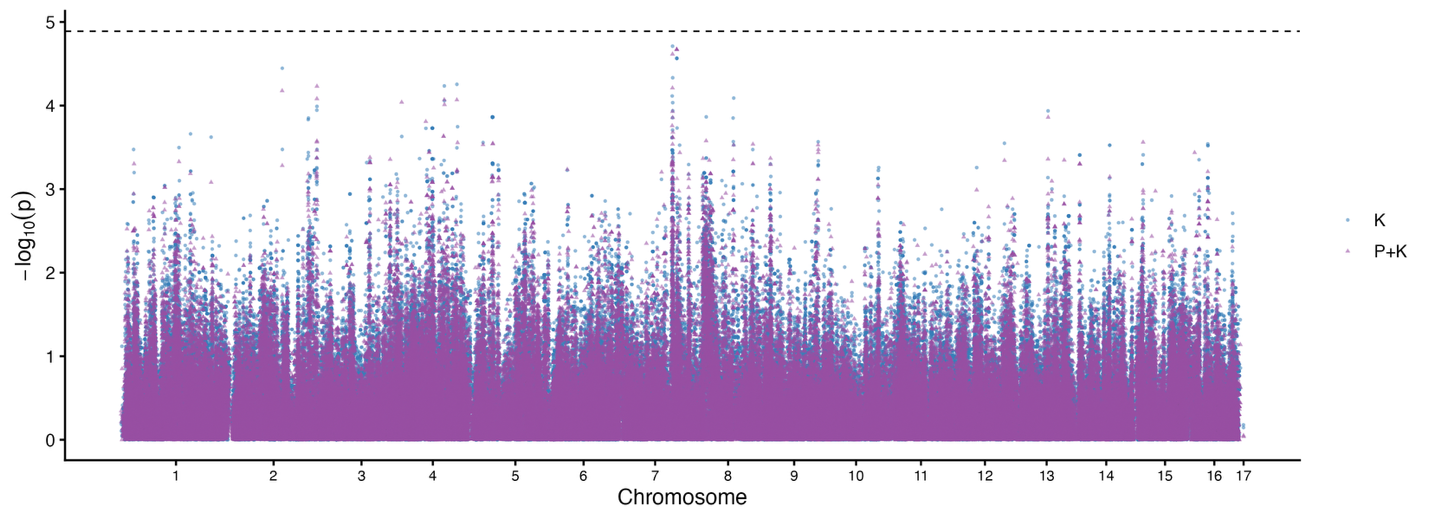

(o)   tip fraction residuals

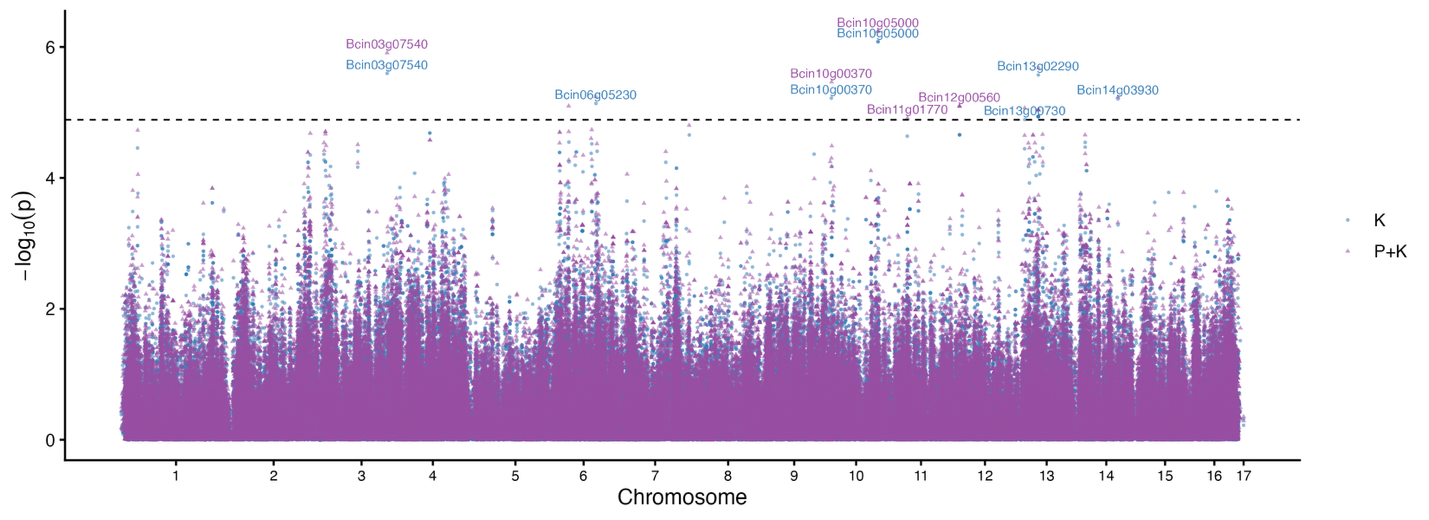

(p) weighted tip fractions

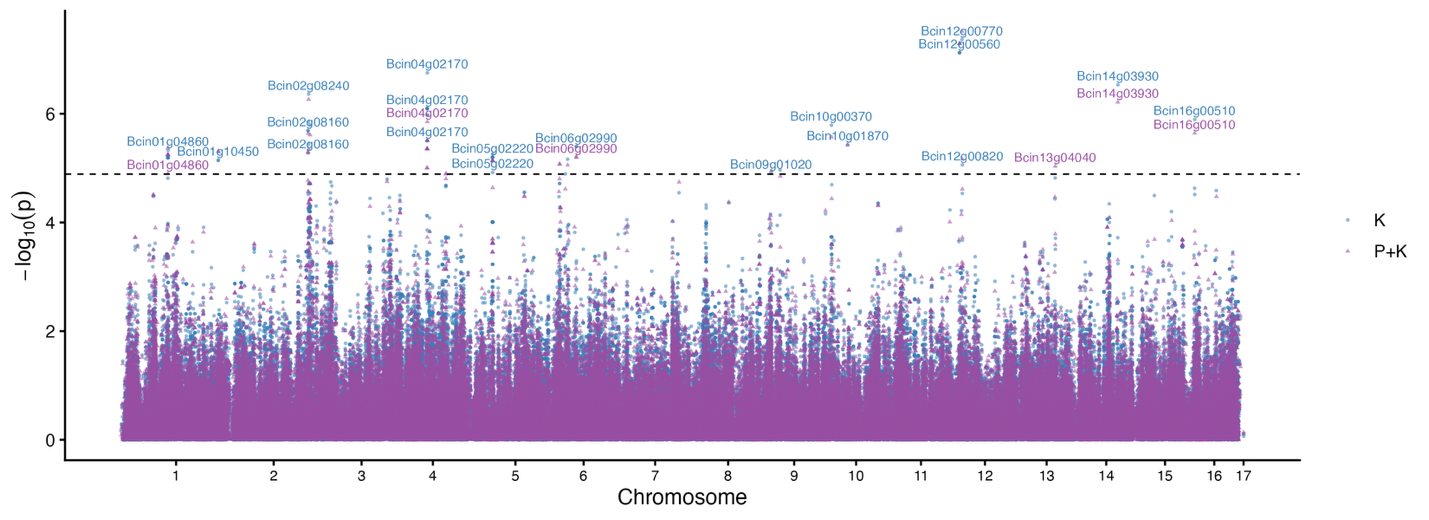

(q) tip density

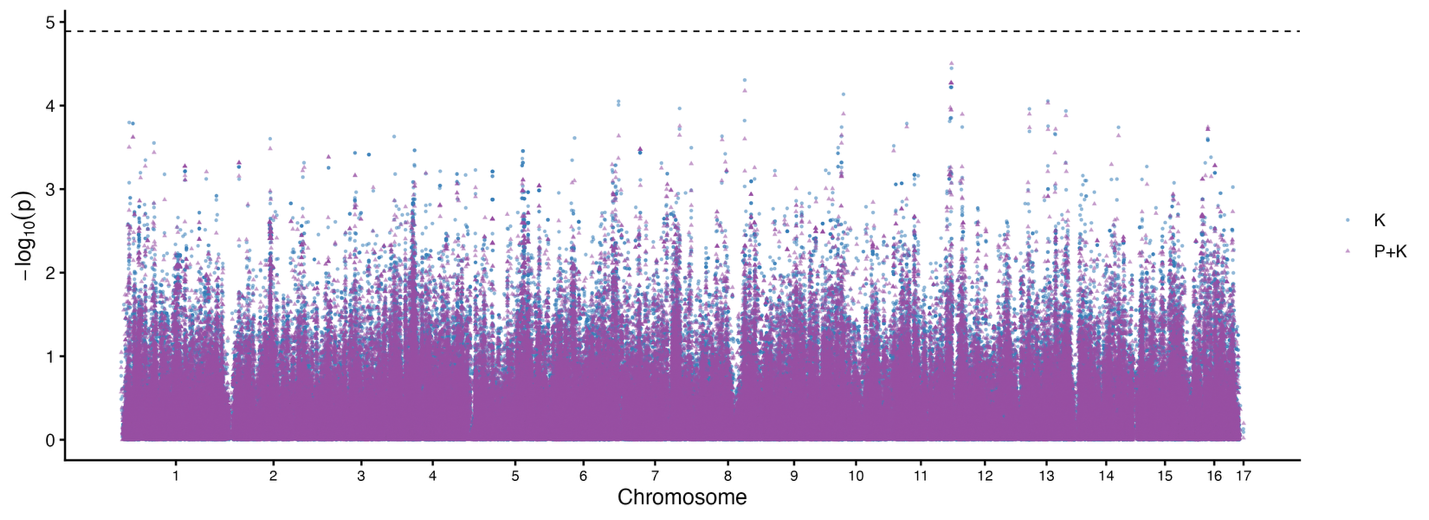

(r) total hyphal length

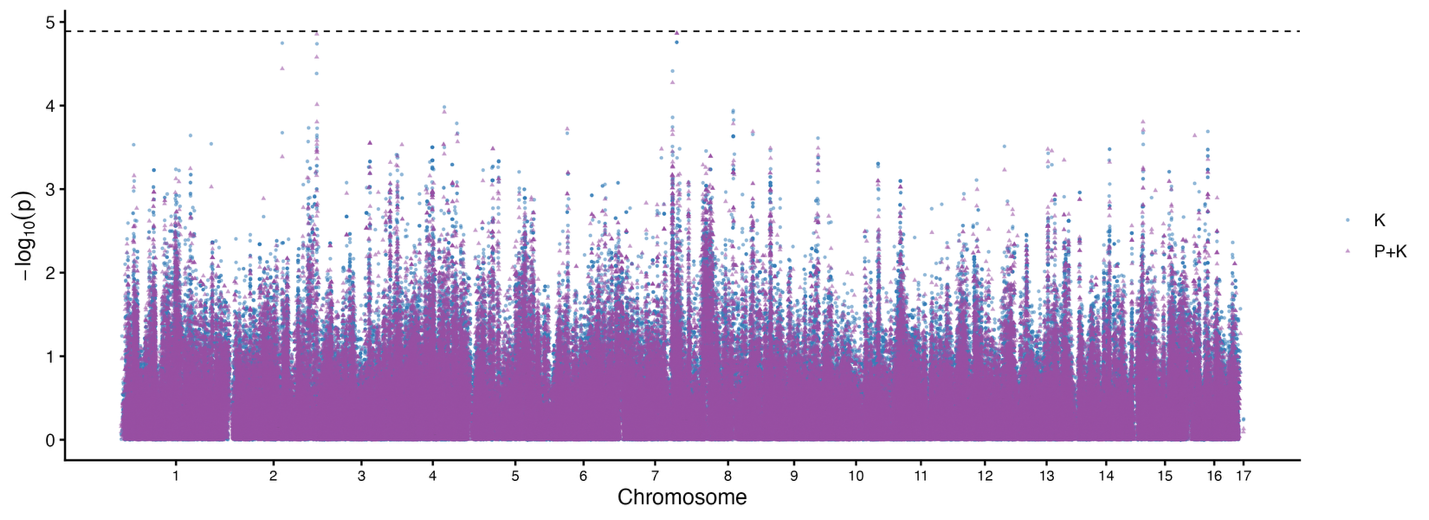

(s) branch fraction

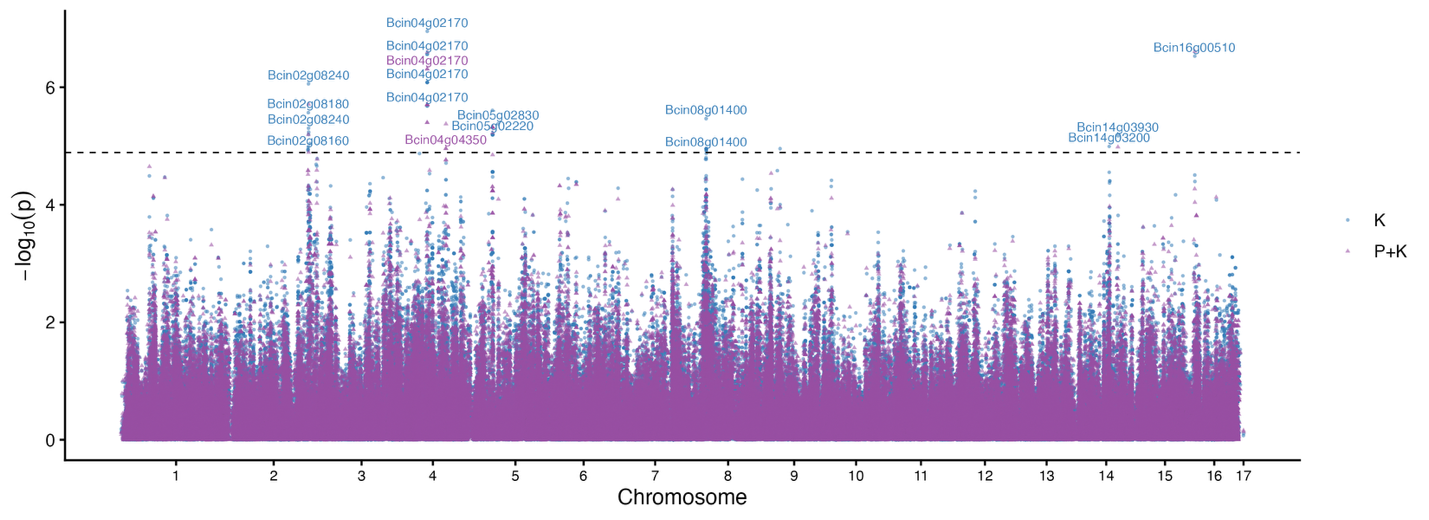

Figure S4. Overlap among significantly associated traits identified from genome-wide association analyses using kinship-corrected models (K) and models incorporating both kinship correction and population structure (P+K). Intersections represent traits sharing significantly associated SNPs across model types, whereas individual set sizes indicate the total number of significant trait associations detected within each model framework.
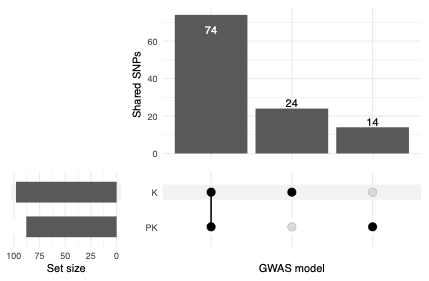
